# ATM is activated by ATP depletion and modulates mitochondrial function through NRF1

**DOI:** 10.1101/414490

**Authors:** Hei-Man Chow, Aifang Cheng, Xuan Song, Mavis R. Swerdel, Ronald P. Hart, Karl Herrup

## Abstract

We have uncovered new insights into the symptoms of ataxia-telangiectasia (A-T). Neurons with high physiological activity, particularly cerebellar Purkinje cells, have large and dynamic ATP demands. Depletion of ATP generates reactive oxygen species that activate ATM (the **A**-**T M**utated gene product). Activated in this way, but not by DNA damage, ATM phosphorylates nuclear respiratory factor-1 (NRF1). This leads to NRF1 dimerization, nuclear translocation and the upregulation of nuclear-encoded mitochondrial genes, thus enhancing the capacity of the electron transport chain (ETC) and restoring mitochondrial function. In cells with ATM deficiency, resting ATP levels are normal, but cells replenish ATP poorly following surges in energy demand and chronic ATP insufficiency endangers cell survival. This is a particular problem for energy-intensive cells such as Purkinje cells, which degenerate in A-T. Our findings thus identify ATM as a guardian of mitochondrial output as well as genomic integrity, and suggest that alternate fuel sources may ameliorate A-T disease symptoms.

**Summary:** Oxidative stress, resulting from neuronal activity and depleted ATP levels, activates ATM, which phosphorylates NRF1, causing nuclear translocation and upregulation of mitochondrial gene expression. In ATM deficiency, ATP levels recover more slowly, particularly in active neurons with high energy demands.

## Introduction

Mitochondrial diseases commonly involve neurological symptoms, and ataxia resulting from cerebellar atrophy and Purkinje cell loss is the most frequent of these (Bargiela et al., 2015). In one cohort study of 345 patients afflicted with a range of different mitochondrial diseases 225 (65%) showed symptoms of ataxia (Bargiela et al., 2015; Lax et al., 2012). The reverse relationship is also found (Bargiela et al., 2015); of persons showing symptoms of definitive ataxia, one-fifth also present with features of mitochondrial dysfunction. Thus, ataxia is linked to mitochondrial defects and vice versa (Fang et al., 2014; Scheibye-Knudsen et al., 2013). This bi-directional correlation led us to consider the protein involved in the inherited ataxia known as ataxiatelangiectasia (A-T), a debilitating autosomal recessive multi-system disease caused by a mutation of the *ATM* gene (Watters, 2003). The protein product of the *ATM* gene was originally identified as a large PI3K-kinase family member that functions as a DNA damage response protein. Yet while the prevalence of many A-T symptoms such as cancer or immunodeficiency can vary, pronounced ataxia, associated with a profound loss of cerebellar Purkinje cells, afflicts all patients with A-T. This is intriguing since, while various mechanisms have been proposed to explain the cerebellar focus of A-T neuropathology, the links between the loss of ATM function and the selective susceptibility of cerebellar neurons to neurodegeneration remain largely unknown.

ATP regulation is critical for a nerve cell. A typical resting neuron contains a billion ATP molecules, yet the firing of only a single action potential is estimated to require the hydrolysis of 10-100 million ATPs to fully restore the resting membrane potential (Howarth et al., 2012; Howarth et al., 2010). This type of estimate underscores the dynamic nature of the ATP supply in neurons and raises questions as to how the levels of such a critical molecule are regulated. Computational modeling predicts that each cerebellar Purkinje cell, with its large cellular size and numerous synaptic connections (Carter and Bean, 2009; Herzig and Shaw, 2018; Sugimori and Llinas, 1990; Welsh et al., 2002), consumes 62 times more energy than a nearby cerebellar granule cell. The majority of this energy is used for the production of action potentials (52%) and the regulation of postsynaptic receptors (38%) with the remainder going towards maintaining membrane potential at rest and other minor tasks (Howarth et al., 2012; Howarth et al., 2010). Thus, neuronal health and survival are heavily dependent on the constant availability of adequate supplies of ATP. Most ATP is derived from glucose through glycolysis (Bratic and Trifunovic, 2010) although the catabolism of other carbon sources may contribute to the process. The predominant site of ATP production is the mitochondrion, through the reactions of the tricarboxylic acid (TCA) cycle and the oxidative phosphorylation (OXPHOS) reactions of the electron transport chain (ETC) (Hall et al., 2012). The TCA cycle in the mitochondrial matrix produces reduced nicotinamide adenine dinucleotide (NADH) and flavin adenine dinucleotide (FADH_2_). These in turn are used as electron donors by the ETC, located in the inner mitochondrial membrane (Owen et al., 2002). The five complexes of the ETC are built from the protein products of literally hundreds of genes, most of which are encoded by the nuclear genome (Dimauro and Rustin, 2009). The highly deleterious effects of mutations in these genes demonstrate that even minor structural changes in ETC proteins disrupt electron transport, ATP production, and can thus cause a range of conditions recognized as mitochondrial diseases that usually have profound impacts on brain functioning.

We report here that a previously unrecognized relationship exists between ATM and the regulation of ATP production in the neuronal mitochondrion. ATM-deficiency results in compromised activities of the TCA cycle and electron transport chain, leading to a reduced capacity to respond to increases in ATP demand. This newly discovered activity of ATM is mediated through nuclear respiratory factor-1 (NRF1). We propose that in the absence of ATM, neurons, in particular mature cerebellar Purkinje cells, cannot respond adequately to the increased in energy demands of neuronal activity. The resulting ATP deficit leads to their degeneration and the observed ataxia and other neurological deficits of A-T.

## Results

### ATM-related deficits in the respiratory chain and TCA cycle

As predicted from the observed correlation between mitochondrial disease and cerebellar ataxia (Bargiela et al., 2015; Lax et al., 2012), the symptoms of A-T cluster with those typically found in diseases involving the mitochondrion (Fang et al., 2014; Scheibye-Knudsen et al., 2013). To confirm this in an unbiased manner, we used the MITODB web application to screen all reported A-T clinical symptoms — neurological as well as peripheral — for their association with mitochondrial function. We found that peripheral symptoms, such as immune deficiency, radiosensitivity, and telangiectasia, failed to show any meaningful mitochondrial association, but CNS phenotypes such as cerebellar atrophy and ataxia showed a strong overlap (Red bars, Fig.1A and Table S1A). This suggests a connection between ATM and mitochondrial function that is most prominent in the nervous system, and is reminiscent of findings from several laboratories, identifying links between mitochondria and A-T (Fang et al., 2016; Patel et al., 2011; Valentin-Vega et al., 2012). Further corroboration comes from the observation that of 8 of the 9 features that are shared by both mitochondrial syndromes and A-T are associated with defects in the ETC (Fig.1B). Taken together, these connections suggested to us that A-T neurodegeneration might be related, at least in part, to defects in OXPHOS and electron transport chain (ETC) function.

**Figure 1.**
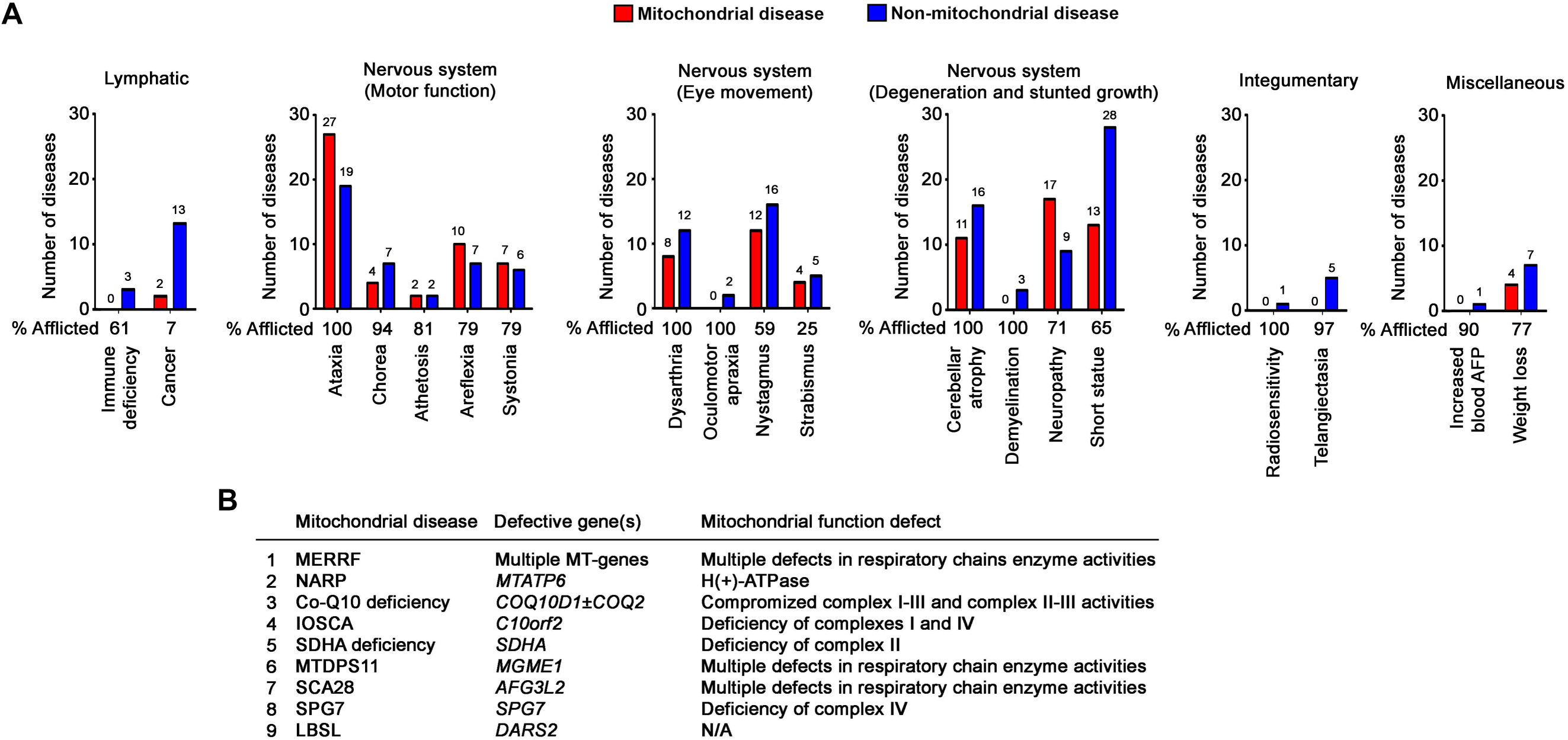
The neurological symptoms from A-T associated with mitochondrial diseases. (A) The number of mitochondrial and non-mitochondrial diseases whose symptoms map to individual clinical symptoms of A-T determined using MitoDB. The percentages of A-T patients associated with each symptom are listed at bottom. (B) A list of nine common mitochondrial diseases associated with symptoms of ataxia and cerebellar atrophy. Eight out of nine are mitochondrial diseases with functional defects in the OXPHOS system.

With this in mind, we re-analyzed earlier microarray results (Li et al., 2013) from human A-T and control cerebellar cortex focusing on genes associated with OXPHOS and ETC functions. Of the more than 31,000 transcripts analyzed, 23% showed significant changes in A-T (Fig.2A, Table S1B). Three-quarters were down-regulated; the rest were up-regulated (Fig.2A, Table S1C). The altered transcripts fell most prominently into 30 Gene Ontology (GO) groups (p<0.0001) as identified by the GATHER platform (Fig.2B, Table S1D) (Chang and Nevins, 2006). Enrichment clustering based on the total number of genes associated with a GO term (Fig.2C) identified several primary metabolic pathways. Using Gene Set Enrichment Analysis (GSEA), we further identified 73 KEGG pathways with nominal *p*-value<0.01 (Table S1E). Pathways associated with OXPHOS (Fig.2D) and the TCA cycle (Fig.2E) were significantly depleted in A-T (adjusted p<0.05).

**Figure 2.**
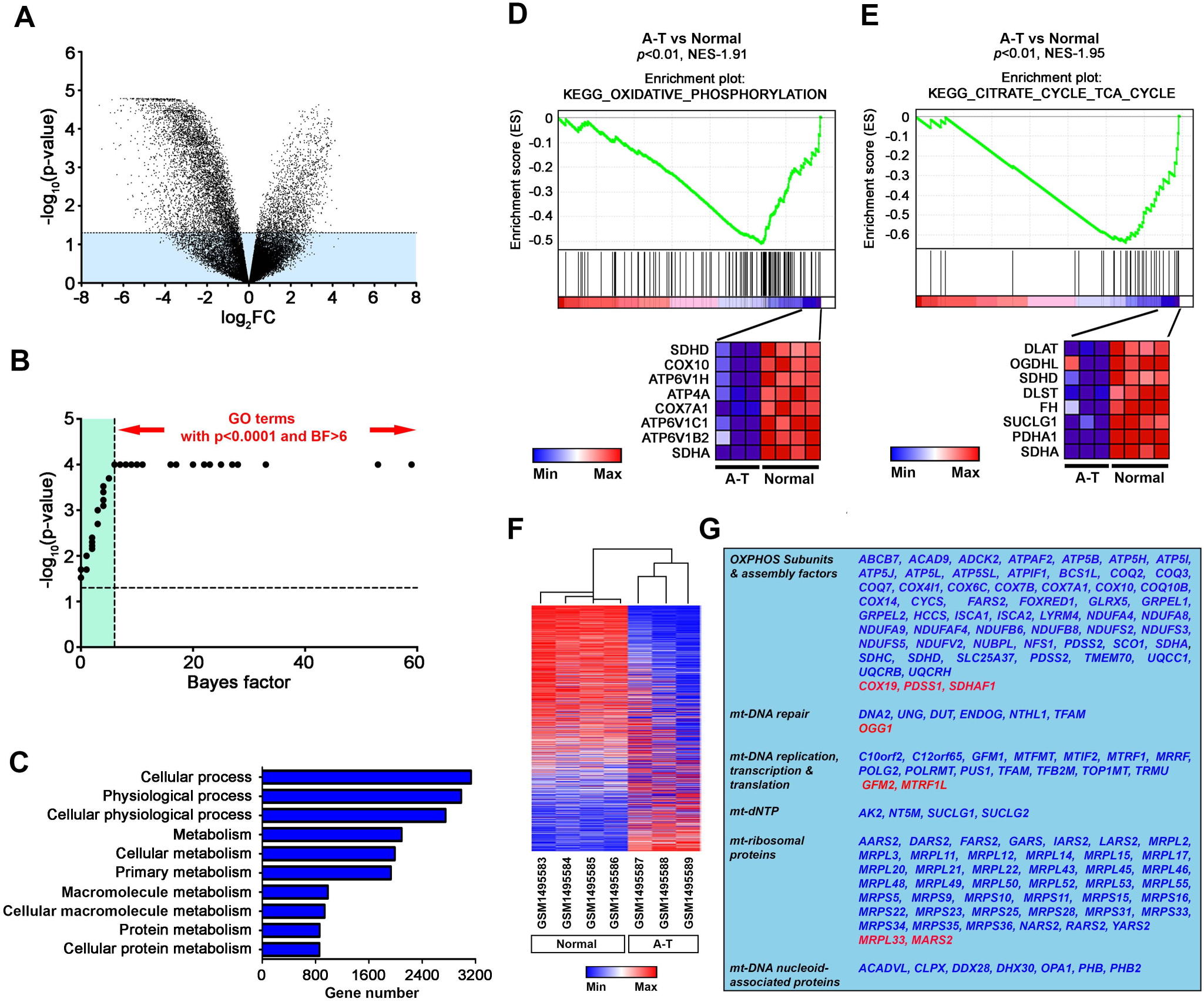
Loss of metabolic function in mitochondria of A-T patients. (A) Volcano plot of differentially expressed genes in cerebellar samples of A-T patients (n=3) versus age-matched controls (n=4). The ‐log_10_[adjusted p-value (p-value)] was plotted against log_2_[Fold Change (FC)]. Genes with significant differences in expression (adjusted p-value<0.05) are in the white area above the dotted line. (B and C) Gene Ontology (GO) analysis of differentially expressed genes (Table S1D). (B) Thirty GO terms with p-value <0.0001 were plotted against the corresponding Bayes factor (BF). Enrichment ranking of pathways with significance metrics above the recommended threshold [P<0.0001 and BF>6] was performed (C). Pathways significantly enriched, determined by the relative number of genes associated within a GO category. (D and E) Gene Set Enrichment Analysis (GSEA) of significantly different genes, among significant differences were pathways of oxidative phosphorylation (D) and the TCA cycle (E). The leading edge (most significant genes), shown as vertical bars, accumulated either left and below the peak of the enrichment score plot (green) or right of the valley, indicating the up‐ or down-regulated genes, respectively. Genes with the highest enrichment score are shown as a gene expression-based heatmap, which indicates relatively reduced expression (blue) of these genes in A-T samples. (F) Heatmap summarizing 1,116 nuclear genes encoding proteins with a high likelihood of mitochondrial localization segregated by function. (G) Significantly changed genes were mostly diminished in A-T (blue gene symbols). Gene symbols marked in red are relatively up-regulated. Note: 42 genes from the original MitoCarta2.0 list failed to match probes from the human expression array.

Together, our in silico analysis suggests that mitochondrial physiology in the cerebellum is significantly altered in the absence of ATM.

Mitochondria are not autonomous entities; they rely heavily on >1,000 imported nuclear-encoded gene products (Lopez et al., 2000). We therefore used the above dataset to perform sub-analyses on 1,158 genes encoding mitochondria-localized proteins as indicated in the MitoCarta 2.0 inventory (Calvo et al., 2016). We observed a distinct pattern of altered expression in these mitochondrial genes in the A-T cerebellar samples (Fig.2F, Table S1F). Expression of 458 (41%) of the nuclear-encoded, mitochondrial genes was significantly altered in A-T. Among these, 83% had reduced expression (p<0.05). The processes that would be affected by the reduced gene expression included virtually every aspect of mitochondrial function – OXPHOS as well as mitochondrial ribosomes, nucleotide levels, mitochondrial DNA repair and replication, and mitochondrial-specific transcription and translation (Fig.2G, Table S1G). This gene regulatory pattern predicts a generalized loss of nuclear-encoded mitochondrial proteins in A-T and suggests that the normal function of ATM is crucial to sustain their presence.

## Reduced expression and function of metabolic enzymes is associated with lack of ATM

Given the conserved nature of most mitochondrial proteins, we sought validation of this prediction in the *Atm*^−/−^ mouse. Biogenesis of a functional OXPHOS system requires not only the core structural proteins of Complex I-V, it also requires an array of OXPHOS assembly factors, as well as proteins needed for coenzyme Q10, cytochrome-c and Fe-S-complex biogenesis (Koopman et al., 2013) (Fig.3A, Table S1H). In the *Atm*^−/−^ mouse cerebellum, as in the A-T human samples (Fig.S1A), expression of genes encoding proteins constituting the OXPHOS core subunits (16/21, p<0.05 – Fig.S1B), OXPHOS assembly factors (17/21, p<0.05 – Fig.S1C), OXPHOS coenzymes and cofactor assembly factors (9/11, p<0.05 – Fig.S1D), mitochondrial ribosomal proteins (8/8, p<0.05 – Fig.S1E); and factors regulating mitochondrial DNA repair, replication, transcription and translation (5/7, p<0.05 – Fig.S1F) were significantly reduced. A notable exception to this downward trend was the increased expression of genes for proteins involved in mitochondrial DNA repair (3/3, p<0.05) and maintenance of the mitochondrial dNTP pool (4/5, p<0.05) (Fig.S1F; Table S1H). The contrary changes in the latter two groups suggests that the cells are mounting a sustained response to enhanced mitochondrial DNA damage. Notably, and consistent with the findings in the *Atm*^−/−^ cerebellum, chronic inhibition of ATM activity in cell culture led to the same expression changes in nuclear-encoded mitochondrial respiratory genes over treatment time (Fig.3B-F). To determine how loss of ATM affected the function of each individual complex, we compared the activities of Complexes I–IV in wild type and *Atm*^−/−^ cultured cortical neurons. All four showed significantly reduced activity (Fig.3G-J, Left panels). The F_1_F_0-_ATP synthase (Complex V) trended lower, but the difference was not significant (Fig.3K, Left panel). This indicates that in the absence of ATM, the functions of the four key ETC complexes are reduced. Importantly, total mtDNA content was not changed in *Atm*^−/−^ neurons (Fig.S1G), implying that the effects seen in the mitochondria are secondary to a defect in nuclear gene expression rather than mitochondrial number. Similar impairment of mitochondrial complex activities was found in wild type neurons with inhibited ATM activity (Fig.3G-K, Right panels). Together, the clinical symptoms, the patterns of gene expression and the reduced functions of the ETC complexes all point to the conclusion that the A-T neurological phenotype has deep roots in neuronal energy metabolism.

**Figure 3.**
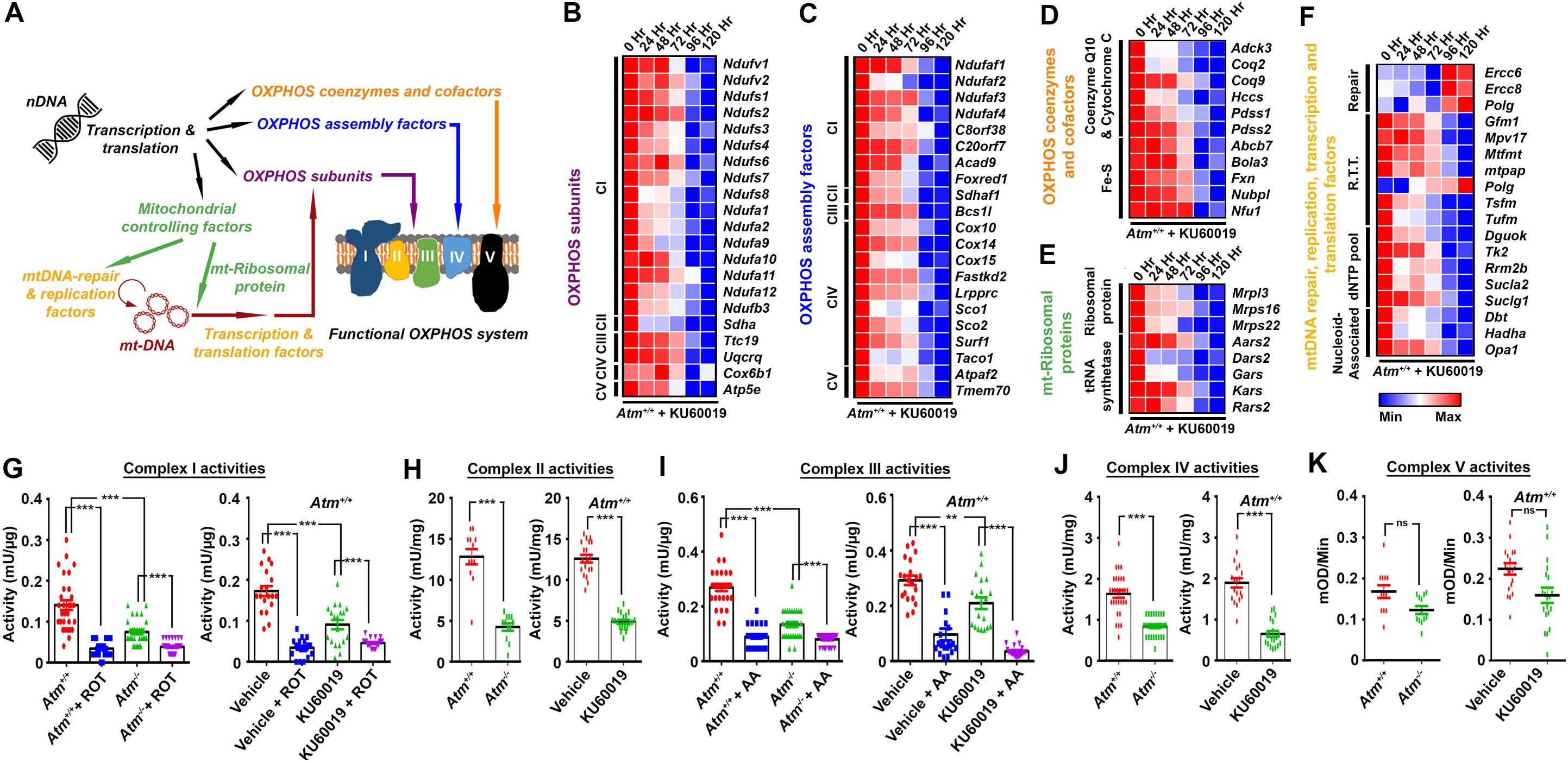
Loss of *Atm* impairs the function of OXPHOS complexes in mouse neurons. (A) Diagram indicating the assembly of the functional OXPHOS system. (B-F) Heatmaps indicating the expression of 79 nuclear-encoded genes whose loss of function results in neurodegeneration in cultured DIV14 primary cortical neurons from wild type mice treated with 1 μM KU60019 for the times indicated (CI through CV labels indicate genes associated with Complexes I through V). (G-K) Activity of each mitochondrial complex measured by colorimetric assays. Left panels: comparisons between complex activities in primary cortical neuron isolated from wild type versus *Atm*^−/−^ mice. Right panels: comparisons between complex activities in wild type cortical neurons treated with vehicle or 1 μM KU60019 for 96 hours. Each group represents the mean±SEM (Complexes I, III, IV and V: N=30; Complex II: N=12) (\*\*\**P<0.001* and *ns*=non-significant, unpaired t-test).

## Altered metabolic dependence of ATM-deficient cells

Previous studies have indicated that TCA cycle metabolites are altered in the cerebellum of *Atm*^−/−^ mice (Fang et al., 2016). We therefore queried our previously published human cerebellar microarray data (Fig.4A) and found that the expression of hexokinase-1/2 (*HK1/2*), the first and rate-limiting enzyme of glycolysis, was significantly reduced in A-T. This suggested that glycolysis might be diminished, and indeed the levels of pyruvate, the end product of glycolysis, were significantly reduced in cultured A-T human fibroblasts (Fig.4B, vehicle condition). We next treated wild type human fibroblasts with 2-deoxyglucose (2-DG), a hexokinase inhibitor (Chen and Gueron, 1992). This caused pyruvate levels in wild type cells to drop to those seen in A-T fibroblasts (Fig.4B). Pyruvate can either enter mitochondria and be converted to acetyl-CoA by pyruvate dehydrogenase (*PDHA1*) or remain in the cytoplasm where it is converted to lactate by lactate dehydrogenase (*LDH*) (McCommis and Finck, 2015). Since the gene expression array data showed that *LDH* was increased and *PDHA1* was dramatically reduced in *Atm*^−/−^ cells (Fig.4A), it appears that pyruvate is both reduced and steered toward the production of lactate in the cytoplasm rather than acetyl CoA in the mitochondrion. Further supporting this hypothesis, UK5099, an inhibitor of the mitochondrial pyruvate carrier [MPC (Hildyard et al., 2005)], resulted in a roughly 50% elevation of pyruvate levels in wild type cells, but no significant change in A-T cells (Fig.4B). This suggests that in ATM-deficient cells pyruvate is converted to lactate in the cytoplasm and thus is unavailable inside mitochondria. Consistent with this idea, treatment with the LDH inhibitor oxamate (Novoa et al., 1959) elevated pyruvate levels in A-T cells while having no significant effect on wild type cells (Fig.4B). Since lactate levels were upregulated by about 3-fold in A-T cells (Fig.4C, vehicle), pCMBS treatment that inhibits lactate efflux from the cell increased lactate inside A-T cells, but had no effect on wild type cells. By contrast, blocking lactate production with oxamate returned lactate in A-T cells to normal levels but had no effect on wild type cells. Mouse primary neurons isolated from *Atm*^−/−^ knockout mice (Fig.S2A) and wild type neurons silenced by *Atm*-specific siRNA (Fig.S2B) both gave similar observations. We next followed the dynamic changes of intracellular pyruvate and lactate in *Atm* knockout neurons over a 24-hour period using FRET nanosensors — Pyronic (PYRuvate Optical Nano Indicator from CECs; San Martin et al., 2014) (Fig.S2C, S2E) and Laconic (LACtate Optical Nano Indicator from CECs; San Martin et al., 2013) (Fig.S2D, S2F). The results with both sensors were consistent with the above results. Thus, in A-T cells mitochondrial pyruvate is lost while cytosolic lactate is increased.

**Figure 4.**
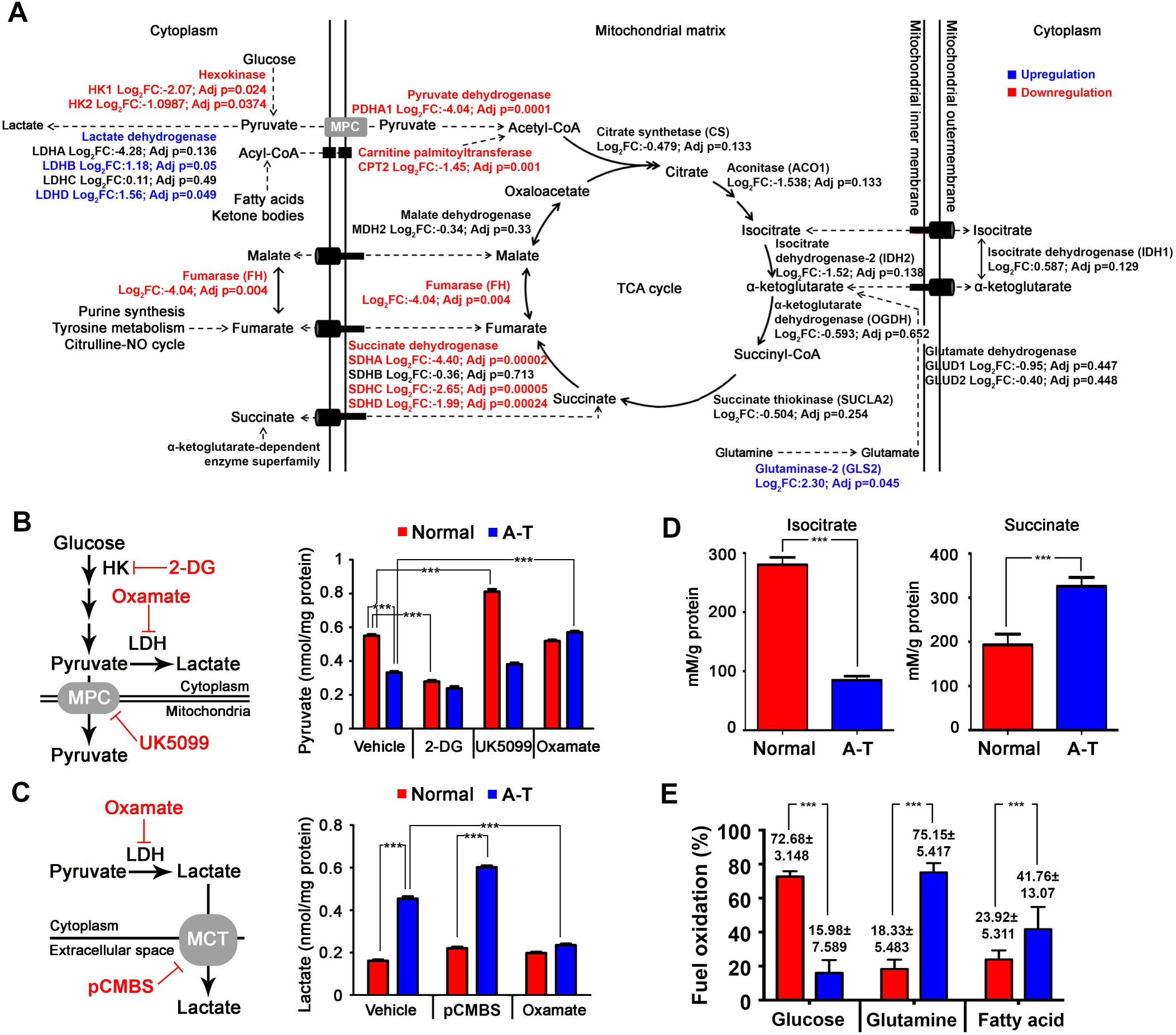
Loss of *Atm* impairs the TCA cycle and favors the conversion of pyruvate to lactate over acetyl-CoA. (A) Diagram of the TCA cycle. Enzymes up-regulated in A-T are shown in blue (adj. p≤0.05); down-regulated enzymes are shown in red (adj. p≤0.05). Those enzymes whose genes had no significant change are shown in black. (B) The levels of pyruvate found after inhibiting glycolysis with 2-deoxyglucose (2-DG) inhibiting mitochondrial import of pyruvate with 1μM UK5099 or inhibiting its cytosolic conversion to lactate with 2.5mM oxamate. The pathways affected by these drugs are diagramed at left (n=18, \*\*\**P<0.001*, One-way ANOVA). (C) The levels of lactate found after inhibition of lactate production by 2.5mM oxamate or blocking release of lactate from the cell by 500 μM pCMBS. Pathways involved are diagrammed at left (n=18, \*\*\**P<0.001*, One-way ANOVA). (D) Changes in levels of isocitrate (left) and succinate (right) in mitochondria of normal or A-T human fibroblast as determined by colorimetric assay (n=18, \*\*\**P<0.001*, *ns=non-significant*, unpaired t-test). (E) Seahorse Mito Fuel Flex test to evaluate cellular respiration of normal and AT human fibroblast in conditions when specific metabolic pathways (glucose, glutamine or fatty acids) were inhibited. The relative glucose, glutamine or fatty acid fuel dependence was calculated. Comparison was made to each fuel source between the normal and A-T groups. See text for details (n=3, ***P<0.001, **P<0.01, ns=non-significant, unpaired t-test).

Once pyruvate enters the mitochondrion it is converted to acetyl-CoA by pyruvate dehydrogenase (PDHA1) before entering the TCA cycle. This conversion would appear to be blocked in A-T cells as there is a significant reduction in PDHA1 (Fig.4A). After this, genes for subsequent steps of the TCA cycle prior to the formation of succinate remained unchanged, but both reactions downstream from succinate – succinate dehydrogenase (SDH) and fumarase (FH) – were significantly reduced (Fig.4A). Consistent with these array results, colorimetric assays revealed that the levels of mitochondrial isocitrate were reduced by more than two-thirds (Fig.4D) while there was a near doubling of mitochondrial succinate levels found in A-T cells (Fig.4E). This situation in A-T cells would seem to be energetically precarious. We therefore considered alternative carbon sources like fatty acid and ketone body catabolism, which generate acyl-CoA and can enter the mitochondria and be converted to acetyl-CoA by carnitine palmitoyltransferase (CPT). Unfortunately, the mitochondrial isoform of *CPT (CPT2)* is reduced by two-thirds in A-T cells (Fig.4A). Earlier findings from our lab showed that dietary supplements of the amino acid glutamine to *Atm*^−/−^ mice alleviated their symptoms and extended their lifespan (Chen et al., 2016a). As the mitochondrial *GLS* isoform, *GLS2*, was increased 5-fold in A-T cells (Fig.4A) it appears possible that A-T cells use glutamine as their source of carbon for the TCA cycle. We confirmed this possibility using the Mito Fuel Flex test. While glucose was the major fuel for wild type cells, A-T cells showed a major dependence on glutamine as an alternative fuel source (Fig.4E).

## ATM-deficient cells have an impaired response to enhanced energy demand

We next investigated how the observed abnormalities affected overall mitochondrial function in ATM-deficient cells. We first used tetramethylrhodamine methylester (TMRM) fluorescence to assess mitochondrial membrane potential (Δψm). In *Atm*^−/−^ neurons, steady state TMRM fluorescence was 1.4-fold higher than in wild type controls (Fig.5A). In examining the cells response to stress, however, an opposite relationship was observed. Oligomycin, an inhibitor of the F_1_F_0_-ATP synthase, caused a rapid reduction in Δψm in *Atm*^−/−^ cells but a slow sustained increase in wild type, presumably because in the presence of oligomycin protons were no longer used to power the ATP-synthase (Fig.5B). The results were similar in human A-T fibroblasts (Fig.S3A-B). Because of the importance of the proton gradient, in cells with impaired OXPHOS Complex V runs in reverse – ATP is hydrolyzed to drive the proton pump and sustain Δψm (Abramov et al., 2010; McKenzie et al., 2007). These observations suggest that the impaired mitochondrial status ATM-deficient cells make them less flexible in adapting to changes in energy demand.

**Figure 5.**
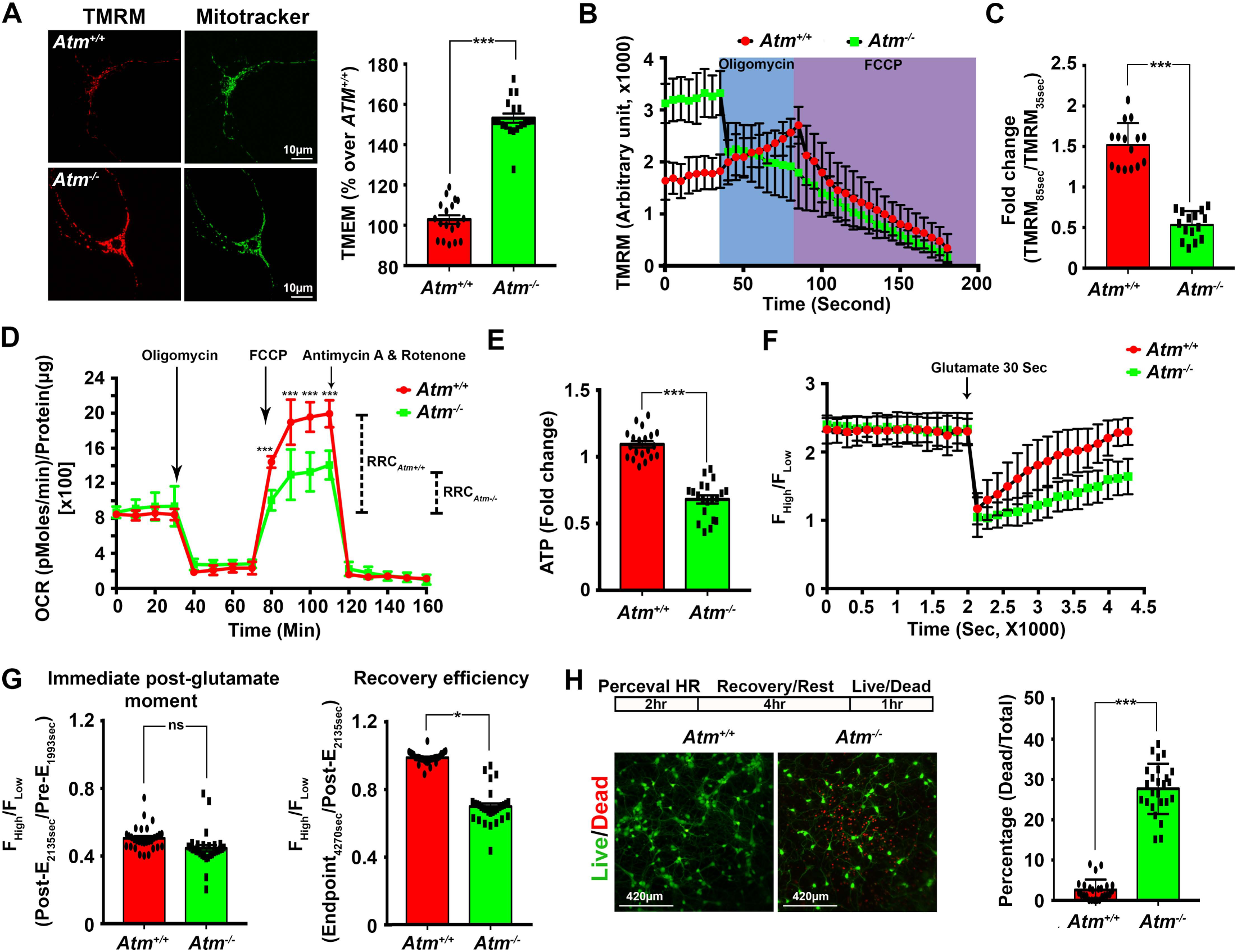
Loss of *Atm* impairs cellular response towards ATP insufficiency in neurons. (A) Static and (B) dynamic measurements of mitochondrial membrane potential in primary cortical neurons evaluated using TMRM, and counterstained with Mitotracker Green (n=15). (C) The relative changes in TMRM signals, expressed as the ratio of OCR values post-oligomycin (85 sec) signal over pre-oligomycin (35 sec; n=15, \*\*\**P<0.001*, unpaired t-test). (D) Seahorse Mito Stress assay of cellular respiration of neurons isolated from *Atm^+/+^* and *Atm*^−/−^ mice (n=16, \*\*\**P<0.001*, two-way ANOVA). (E-F) Static (E) and dynamic (F) changes in ATP levels in wild type and *Atm*^−/−^ primary cortical neurons measured by the Perceval HR reporter. (E) Equilibrium ATP levels at the instant prior to the dynamic experiment (n=20, *\*\*\*P<0.001*, unpaired t-test). (F) Live-imaging of cultured neuron responses to synaptic activity-dependent energy consumption. After establishing a steady baseline, 50 μM glutamate was added for 30 seconds (arrow) to excite the neurons and cause ATP to be consumed; glutamate was then washed away and intracellular ATP levels allowed to recover (n=30, *\*\*\*P<0.001*, two-way ANOVA). (G) The relative abundance of ATP immediately following the removal of glutamate (Signal ratio: Post-E_2135sec_/Pre-E_1993sec_, left panel) and after recovery (Signal ratio: Endpoint_4270sec_/Post-E_2135sec_, right panel) (n=30, *\*\*\*P<0.05*, *ns*=non-significant, unpaired t-test). (H) Representative images of the live/dead status of neurons four hours after Perceval HR assay. (Live cell: Green; Dead cell: red). Quantification shown in right panel (n=25, *\*\*\*P<0.001*, unpaired t-test).

To validate this suggestion, we measured the dynamic changes in oxygen consumption rate (OCR) – an indicator of mitochondrial respiration. We found that the basal OCR in mouse *Atm*^−/−^ neurons (Fig.5D) and human A-T fibroblasts (Fig.S3D) was not significantly different from controls. The drop in OCR after treatment with oligomycin reveals the oxygen consumed by the ATP synthase. The nearly identical response of A-T and control cells to the drug (Fig.5D, Fig.S3D) demonstrates that under resting conditions, the respiratory activity of both wild type and ATM-deficient cells are similar. However, subsequent treatment of the cells with FCCP (a weak lipid soluble acid that allows proton leakage across the mitochondrial membrane and thus drives maximal OCR) revealed that ATM-deficient cells had a reduced response compared to wild type. This in turn implies that the reserve respiratory capacity (RRC) of ATM deficient cells is significantly smaller than wild type controls (Fig.5D, S3D; dashed lines). This reduced RRC, together with our data showing that ATM deficient cells hydrolyze ATP to maintain their Δψm, lead to the inevitable conclusion that less ATP is available in response to surges in neuronal energy demand. Indeed, under equilibrium conditions, we found that, compared to controls, the levels of ATP were reduced by one quarter in mouse *Atm*^−/−^ cortical neurons (Fig.5E) and by nearly one half in A-T human fibroblasts (Fig.S3E). We synaptically activated the cultured neurons for 30 seconds with glutamate to produce large fluxes of sodium and potassium and thus stimulate the accelerated hydrolysis of ATP by the Na^+^/K^+^-pump. We then monitored the ATP:ADP ratio in real time using the PercevalHR biosensor (Tantama et al., 2013). In wild type cultures, glutamate treatment resulted in a rapid 50% reduction in the ATP:ADP ratio but over the next 30 minutes the ratio returned to pre-stimulus levels (Fig.5F-G). In *Atm*^−/−^ cultures, while the initial reduction in the ATP:ADP ratio was nearly the same as in controls, the rate of recovery was significantly slower (Fig.5F-G). Emphasizing this point further, while wild type cells could recover fully after glutamate, the treatment killed nearly a quarter of the *Atm*^−/−^ cells (Fig.5H). These data show that cells lacking ATM are unable to respond to increased energy demands and are thus more vulnerable during periods of high activity.

## Nuclear respiratory factor-1 links ATM activity and mitochondrial function

To determine how ATM deficiency led to the observed defects in ATP generation, we returned to the global gene expression data from human cerebellum (Fig.2A). We analyzed all possible transcript isoforms of the genes that were differentially expressed in A-T and control samples (12,931 transcripts from 7,222 genes). ATM itself is a serine/threonine kinase with no known DNA binding capacity; if it were to modulate gene expression it would most likely work through modulating activities of one or more transcription factors. We therefore searched the promoter regions (−450 to +50 bp) of all differentially-regulated transcripts for transcription factor binding sequences using the PSCAN tool (Zambelli et al., 2009). This analysis identified 71 candidate transcription factors that were significantly associated with the differentially expressed genes (p<0.05, Z-test; Table S1J). Of these 71, binding motifs for NRF1 (nuclear regulatory factor-1) ranked highest (*p*=8.24 × 10^−114^; Fig.6A). Among the 12,931 transcripts, binding sequences with >85% similarity to the NRF1 consensus motif were found in about one-third of them (4,624 promoters belonging to 3,360 genes, Fig.6B-C). NRF1 is a basic leucine zipper (bZIP) transcription factor (Fig.6B) that functions as a trans-activator of nuclear-encoded respiratory genes (Evans and Scarpulla, 1990; Scarpulla, 2008) as well as performing extra-mitochondrial regulatory roles (Satoh et al., 2013). With its emergence as a strong candidate to explain the ATM-mediated changes in ATP, we analyzed the expression patterns of the 3,360 genes identified as potentially regulated by NRF1 (Table S1K) and found that over 80% of them were significantly reduced in AT cerebellar samples (Fig.6D, Table S1L). Further, of 2,470 NRF1 targets identified in both the ENCODE database and a previous NRF1 ChIP-Seq study (Satoh et al., 2013) we found that 440 appeared in our list of genes with altered expression in A-T (Fig.6E). KEGG pathway analysis of the 440 common NRF1 targets from the A-T regulated genes identified cell cycle, Wnt signaling, oxidative phosphorylation and circadian rhythm pathways as significantly affected (Fig.6F). We next used immunohistochemistry to determine the anatomic location of the NRF1 protein in normal and ATM-deficient brain. In wild-type mouse and control human samples, we found that the NRF1 immunoreactivity was substantially enriched in Purkinje cells compared to other cerebellar cell types. In the absence of ATM, NRF1 immunostaining in Purkinje cells was reduced and this reduction was most dramatic in the cell nucleus (Fig.6G). The implication of these findings is that ATM regulates NRF1 levels as well as its nuclear-cytoplasmic shuttling.

**Figure 6.**
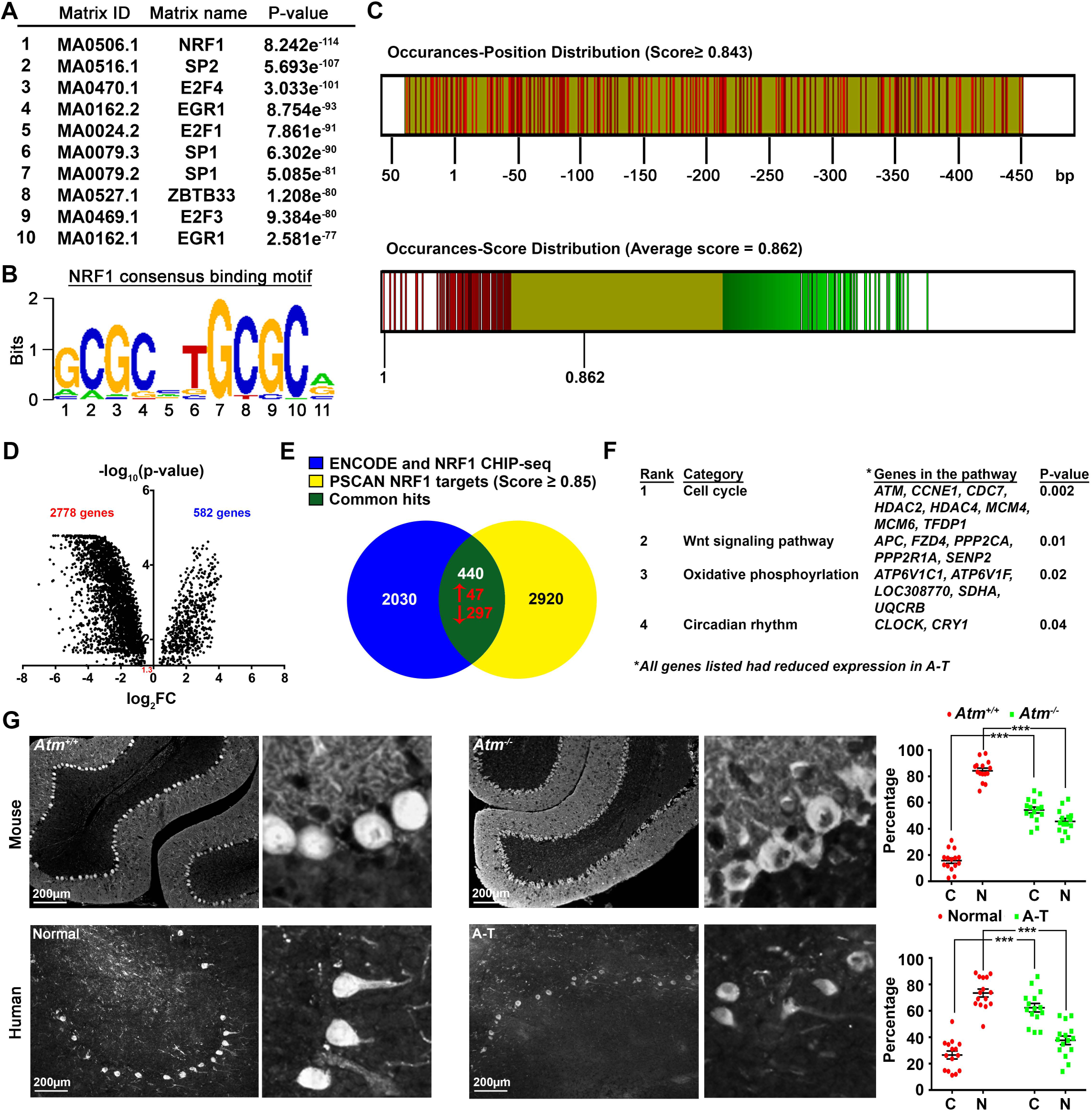
Nuclear respiratory factor-1 (NRF1) links ATM activity and mitochondrial function. (A) The top 10 over-represented transcription factors predicted by the PSCAN platform to bind promoters of nuclear-encoded mitochondrial genes, ranked according to their z-test P-value. (B) The consensus NRF1 binding site for these transcription factors. (C) Top panel shows the location of the best predicted NRF1 binding motifs within the −450 to +50 bp region on the tested promoters with corresponding matching scores (the value representing the likelihood for NRF1 to bind to the promoter) higher than the average genome-wide matching score 0.843. Bottom panel shows the distribution scores for tested promoters, with an average matching score values of all predictions equaling 0.862, higher than the average genome-wide matching score 0.843. These, together with the z-test P-value shown in Fig.6A (NRF1 = 8.242 × 10^−114^) imply that the NRF1 binding motif is non-randomly enriched in the tested set of promoters, in which the gene expression level was significantly altered in A-T cerebella samples. Predictions on each promoter were colored as straight lines according to their corresponding matching score (red=high; yellow=medium; green=low). (D) Volcano plot showing the p-value of the relative expression levels of the genes predicted to be regulated by NRF1. (E) Venn diagram illustrating the overlap between the set of genes computationally identified as NRF1 targets (yellow) and the combined sets of observed NRF1 targets found in the ENCODE database and a published NRF1-Chip sequencing study (blue). (F) A table showing the four KEGG pathways identified using genes whose promoters scored as one of the 400 top hits for likelihood as an NRF1 promoter. Genes associated in the pathways as well as the corresponding p-values are shown. (G) Representative images of NRF1 immunostaining in mouse (upper panels) and human (lower panels) cerebellar tissue samples from wild type (left) and ATM-deficient (right) subjects. Quantification of Purkinje cells with cytoplasmic (C) and nuclear (N) NRF1 in cells are shown on the right (n=15, *\*\*\*P<0.001,* one-way ANOVA).

## ATP depletion activates ATM with downstream effects on NRF1

Several studies have shown that ATM can function as a redox sensor because it can be activated by oxidative stress independently of its response to DNA damage (Ashley et al., 2012; Guo et al., 2010). We wondered, therefore, whether transient reductions in ATP might lead to ETC imbalances and that the resulting increase in reactive oxygen species (ROS) might activate ATM. To test this, we induced a modest depletion of ATP by treating cells with atractyloside (an inhibitor of the ADP/ATP carrier) or a more dramatic ATP depletion with oligomycin. Oligomycin resulted in significant cell death due to the severe ATP depletion (Fig.S4A-B); yet neither atractyloside nor oligomycin induced a significant DNA damage response (Fig.7A). Confident that our results would not be confounded by an independent response of ATM to DNA damage, we used the less toxic atractyloside to test primary cultures of mouse neurons. We observed a good correlation between the atractyloside-induced oxidative stress and its progressive depletion of ATP, consistent with our hypothesis that cellular ATP depletion induces oxidative stress (Fig.S4C). To determine whether this ATP depletion-induced oxidative stress could activate ATM and impact NRF1, we tracked the subcellular location of NRF1 and activated (phosphorylated) ATM. Oxidative damage induced either directly with hydrogen peroxide (H_2_O_2_) or indirectly with atractyloside resulted in the nuclear localization of both phospho-ATM and NRF1 (Fig.7B-C). Significantly, the actions of both drugs were blocked by the anti-oxidant, N-acetyl cysteine (NAC) (Fig.7B-C). We determined that ATM was required for this response as NRF1 nuclear localization in response to either H_2_O_2_ or atractyloside was lost in *Atm*^−/−^ neurons (Fig.7B-C). Equally important, induction of DNA damage alone was insufficient to trigger the NRF1 response. Etoposide, a topoisomerase II inhibitor that leads to DNA double strand breaks (Fig.7A), failed to reduce ATP levels (Fig.S4A) or stimulate the translocation of NRF1 to the nucleus (Fig.7B-C).

**Figure 7.**
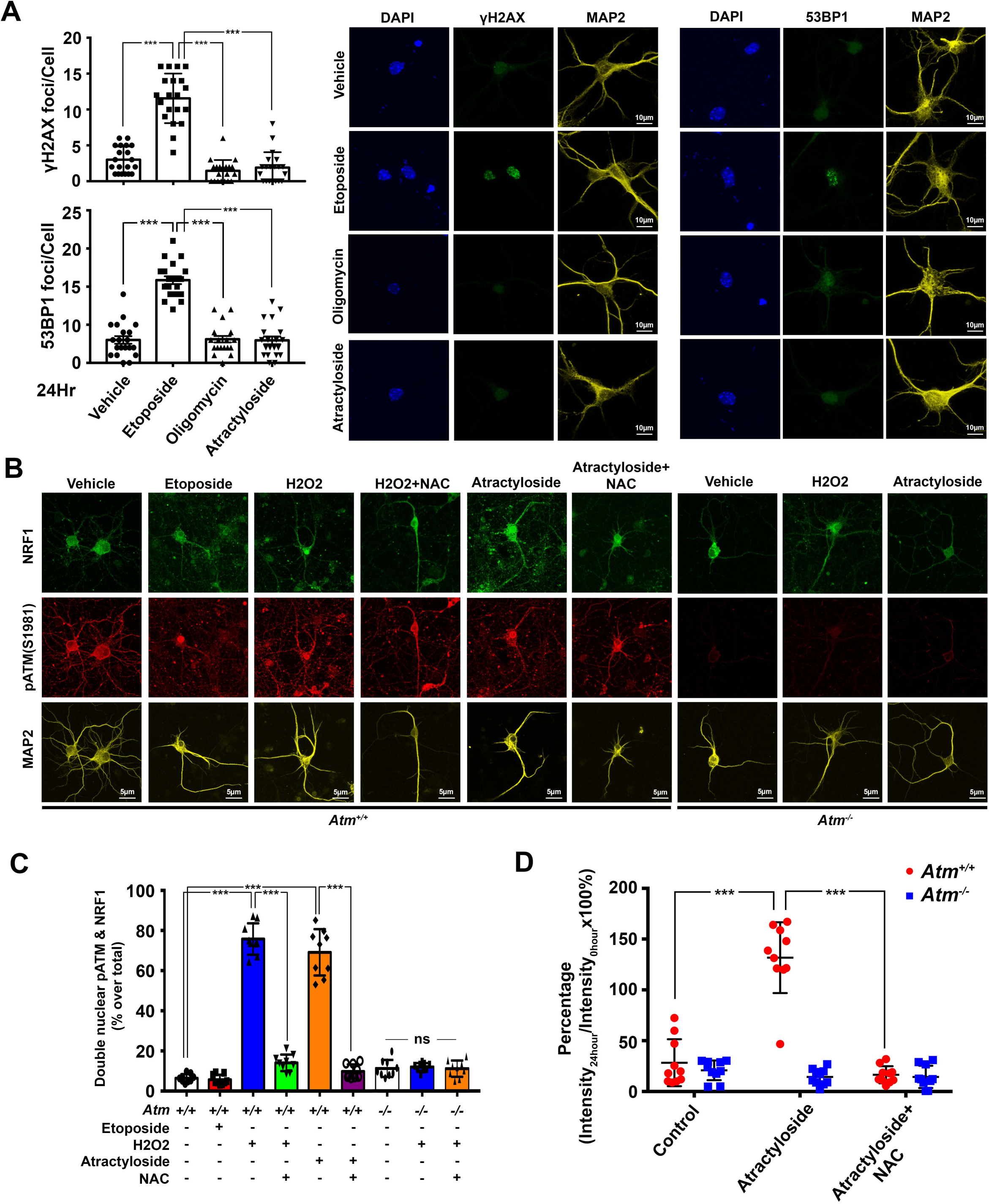
ATP depletion activates ATM with downstream effects on NRF1. (A) DIV14 Primary cortical neurons isolated from wild type mice were treated with vehicle, 1 μM etoposide, 0.5 μM oligomycin or 1 μM atractyloside for 24 hours. The number of H2AX and 53BP1 foci in MAP2^+^ neurons were then quantified (n=20, *\*\*\*P<0.001*, one-way ANOVA). (B) Immunocytochemistry for activated ATM [pATM(S1981)], NRF1 and MAP2 (as indicated) in wild type (*Atm*^*+/+*^ – left) and *Atm*^−/−^ (right) DIV14 primary cortical neurons treated with either vehicle, 1 μM etoposide, 10 nM H_2_O_2,_ or 1 μM atractyloside in the presence or absence of 5 mM N-acetylcysteine (NAC) for 24 hours. (C) Quantification of the results in panel B (n=9, *\*\*\*P<0.001*, *ns*=non-significant, one-way ANOVA). (D) NRF1-mediated transcription activities were measured using the NRF1mitoGFP reporter in wild type DIV14 primary cortical neurons treated with either vehicle or 1μM atractyloside in the presence or absence of 5mM NAC. Intracellular GFP signals resulting from NRF1 transcription activity were monitored for 24 hours after initiation of drug treatments. The ratios of cellular GFP intensities at 24-hour over 0-hour time points are shown. Representative images of the reporter signals at 0- and 24-hour time points can be found in Fig.S4D.

NRF1 helps to coordinate the expression of nuclear and mitochondrial genomes. We therefore suspected that nuclear localization of NRF1 was predictive of its transcriptional activity. We tested this using the live-cell reporter, NRF1mitoGFP, which contains an NRF1 binding sequence driving GFP protein expression fused with a mitochondrial localization signal. This allowed us to simultaneously monitor NRF1-mediated transcriptional activity as well as mitochondrial morphology (Hodneland Nilsson et al., 2015). As predicted from the NRF1 nuclear localization, we observed that moderate but chronic depletion of ATP induced NRF1-mediated transcription (Fig.7D and S4D). In contrast, when ATM activity was lost, or with NAC co-treatment, induction of NRF1 activity was abolished (Fig.7D and S4D). Significantly, in the presence of ATM, the mitochondria appeared to maintain their normal filamentous morphologies while in ATM-deficient situations, the mitochondria appeared more fragmented (Fig.S4E). Thus, in conditions when cellular ATP is chronically in demand, ATM is required to stimulate NRF1 activity and upregulate mitochondrial capacities. These results suggest that the oxidative activation of ATM induced by suboptimal cellular ATP levels results in nuclear localization of NRF1 with important consequences for nuclear encoded mitochondrial gene expression. The strong implication of the data is that ATM is a transducer of deficits in the energy status of a cell.

We next asked how the kinase activity of ATM was able to alter the transcriptional activity of NRF1. NRF1 binds to its DNA recognition site as a homodimer whose formation is triggered by phosphorylation of the monomer (Gugneja and Scarpulla, 1997). While *NRF1* gene transcription levels were not significantly different (Fig.8A), Western blots revealed that in ATM-deficient cells the level of NRF1 protein was reduced and the fraction of total NRF1 that was present as a dimer dropped significantly (Fig.8B). Scrutiny of the NRF1 protein sequences of four mammalian species identified a total of eight well-conserved (S/T)Q sequences – canonical ATM target sites (Fig.8C). To determine whether these sites were ATM-targeted we co-expressed FLAG-tagged wild type (WT) ATM and GFP-tagged NRF1 in *Atm*^−/−^ in mouse embryonic fibroblasts. Subsequent immunoprecipitation revealed a robust physical interaction between the two proteins (Fig.8D). Further, Western blot of the immunoprecipitates probed with a phospho(S/T)Q antibody revealed a strong band that was missing if a kinase-dead (KD) FLAG-ATM construct was used instead of a wild type one (Fig.8E). (S/T)Q phosphorylation of NRF1 was also lost in cultures treated with an ATM inhibitor (1 μM KU60019) but not with inhibitors of ATR (0.5 μM VE821) or DNA-PK (20 nM NU7441) (Fig.8F). These data suggest that specific ATM phosphorylation facilitates the homodimerization of NRF1. Although there are a total of eight potential (S/T)Q phosphorylation sites on NRF1 (Fig.8C), only the T259 residue located within the DNA-binding and dimerization (DIM) domain was previously identified by mass spectroscopy (Hornbeck et al., 2015) as a likely site for the dimerization site. In our search for the relevant ATM target we therefore focused on T259 and created mutant clones of the NRF1 protein (Fig.8G). In the presence of wild type ATM, co-expression of the wild type NRF1 protein or the NLS-deletion mutant (NLSdel) resulted in robust phosphorylation of the (S/T)Q site (Fig.8G). However, when the deletion was extended into the DNA-binding and dimerization domain (DIM) (DIMdel: T87-E283), the (S/T)Q phosphorylation signal was lost (Fig.8G). Refining the ATM target residue still further, expression clones that carried threonine-to-alanine triple (T201A, T259A and T287A), double (T201A and T259A) mutations, or the single (T259A) mutation resulted in similar loss of (S/T)Q phosphorylation (Fig.8G), indicating these residues are potential ATM targets. Testing single point mutant clones at each of these sites confirmed the T259 residue as the relevant target of the ATM kinase (Fig.S5A).

**Figure 8.**
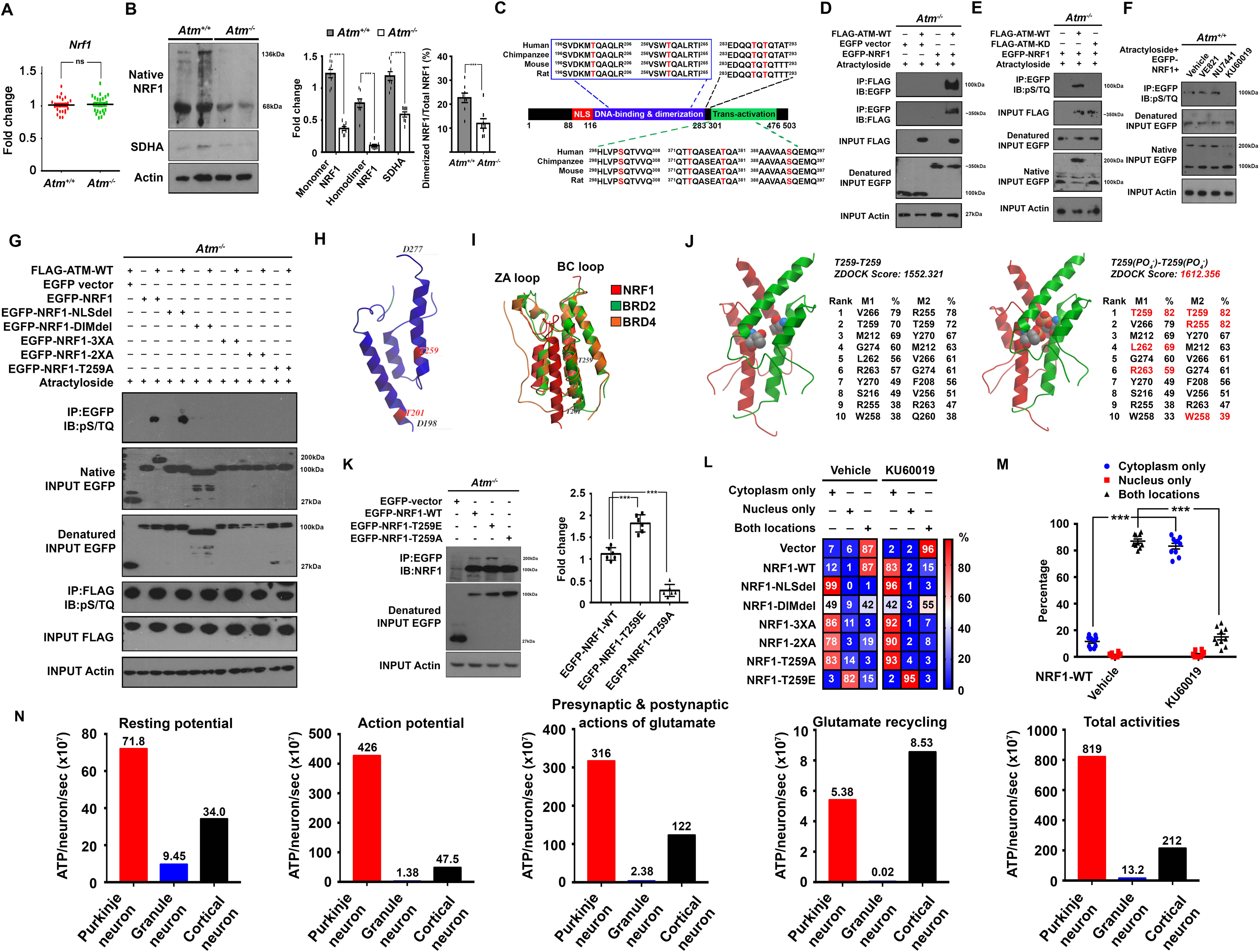
ATM-mediated phosphorylation of the NRF1 T259 residue facilitates homodimerization and nuclear localization. (A) *NRF1* gene expression in mouse cerebellum (n=28-30, ns = not significant, unpaired t-test). (B) Representative Western blot (left panel) and quantification (right panel) of monomeric and dimeric NRF1 and one of its target, succinate dehydrogenase, in mouse cerebellum (n=4, *\*\*\*P<0.001*, unpaired t-test). (C) NRF1 protein sequence alignment, performed on the Clustal Omega platform. (D-F) Representative Western blots of immunoprecipitates (IPs) showing interactions between NRF1 and ATM. (D) Reciprocal IPs of EGFP-NRF1 and FLAG-ATM-WT; (E) (S/T)Q phosphorylation (top) and dimerization of EGFP-NRF1 mediated by wild type (WT) but not kinase dead (KD) ATM (F) Inhibition of ATM (with KU60019), but not ATR (with VE821) or DNAPKcs (with NU7441), blocks (S/T)Q phosphorylation of EGFP-NRF1 in mouse embryonic fibroblasts. (G) Western blot images showing that both the nuclear localization of NRF1 and its phosphorylation on T259 are required for NRF1 homodimerization (n=4). (H) NRF1 (aa198-271) structure is predicted to adopt a bromodomain configuration in the region of its DNA-binding and dimerization domain. (I) This region of NRF1 (red) superimposes well with BRD2 and BRD4 bromodomains (green and orange respectively). (J) *in silico* site-directed mutagenesis was performed to obtain the T259(PO4^−^) structure followed by homodimer docking analyses using the ZDOCK platform. Predicted homodimerized best-ranking NRF1 bromodomain structures are shown in the table. Contact area analysis was performed using the ICM-browser pro. Amino acid residues that showed significant interactions between the monomers were ranked according to the percentage of contacting area over total exposed area. (K) Western blots of GFP tagged-NRF1 co-immunoprecipitations. Quantification of the percentage of dimerized NRF1 is shown (n=6, *\*\*\*P<0.001*, one-way ANOVA). (L) Nuclear versus cytoplasmic localization of NRF1 mutants expressed in in DIV12 primary cortical neurons as shown in a heatmap. (M) Distribution of EGFPNRF1-WT proteins in neurons treated with 1 μM KU60019 for 96 hours (n=10, *\*\*\*P<0.001*, one-way *ANOVA*). (N) Histograms showing ATM requirements of individual Purkinje (red), cerebellar granule (blue) or cortical pyramidal (black) neurons. Individual panels show energy consumption by specific subcellular processes as marked. Parameters and calculations as per Howarth et al (2012), details can be found in Table S1O.

We further explored the nature of NRF1 dimerization *in silico*. While the structural features of the entire NRF1 protein have not yet been identified, we used *ICM-Pro Browser* to superimpose the *Phyre*^*2*^-predicted structure of the NRF1 dimerization domain (DIM) onto the bromodomain-containing protein-2/4 (BRD2/4 – Garcia-Gutierrez et al., 2012). The fit was good, suggesting that the DIM domain of NRF1 closely resembles a bromodomain (Fig.8H-I). *In silico* phosphorylation of T259 followed by *ZDOCK* docking and contact area analyses with *ICM-Pro Browser* suggested a more favorable interaction between monomers phosphorylated at T259 than between nonphosphorylated monomers (Fig.8J, Table S1M, N), consistent with the idea that T259 phosphorylation is a post-translational modification that triggers homodimerization (Fig.8K, S5B). We separately verified the impact of NRF1 phosphorylation on nuclear-cytoplasmic shuttling (Fig.8L-M). Expression of a phosphomimetic mutant of NRF1 (T259E) promoted its nuclear localization while expression of non-phosphorylatable NRF1 (T259A) led to a more prominent cytoplasmic localization in primary neurons. As expected, nuclear localization of wild type NRF1 was lost if cells were pre-treated with ATM inhibitor (Fig.8L-M). We also validated the importance of ATM activity (Fig.S5C-D) – specifically the activity induced by oxidative stress (Fig.S5E) – as mediating the phosphorylation, dimerization and nuclear localization of NRF1.

## Ectopic expression of T259E NRF1 rescues bioenergetic defects in *Atm*^−/−^ neurons

To confirm that NRF1 is the downstream effector of ATM in sustaining mitochondrial bioenergetics, and that the NRF1 T259 phosphorylation site plays an indispensable role in mediating mitochondrial dysfunction and ATP insufficiency in A-T, we ectopically expressed either the phosphomimetic T259E or non-phosphorylatable T259A mutants of NRF1 in neurons for 96 hours prior to an array of mitochondrial analyses. Real-time PCR revealed that NRF1-T259E but not T259A rescued the reduced mRNA expression of most nuclear-encoded mitochondrial respiratory genes in *Atm*^−/−^ cells (Fig.9A). This rescue effect was subsequently reflected in multiple mitochondrial functions, including the static (Fig.9B-C) and dynamic (Fig.9D-E) mitochondrial potential; the reserved respiration capacity (Fig.9F); the static levels of ATP (Fig.9G); and the dynamic ATP recovery rate (Fig.9H-J). None of the restorative effects of the NRF1-T259E mutant was seen with the non-phosphorylatable NRF1-T259A mutant (Fig.9K). Conversely, using *Atm*^*+/+*^ neurons, we confirmed that NRF1 nuclear activities depend on ATM activity (Fig.9L-M), mediated by specific phosphorylation of the T259 residue, even in the presence of functional ATM, presumably due to a dominant-negative activity from overexpression of non-phosphorylatable T259A (Fig.9N).

**Figure 9.**
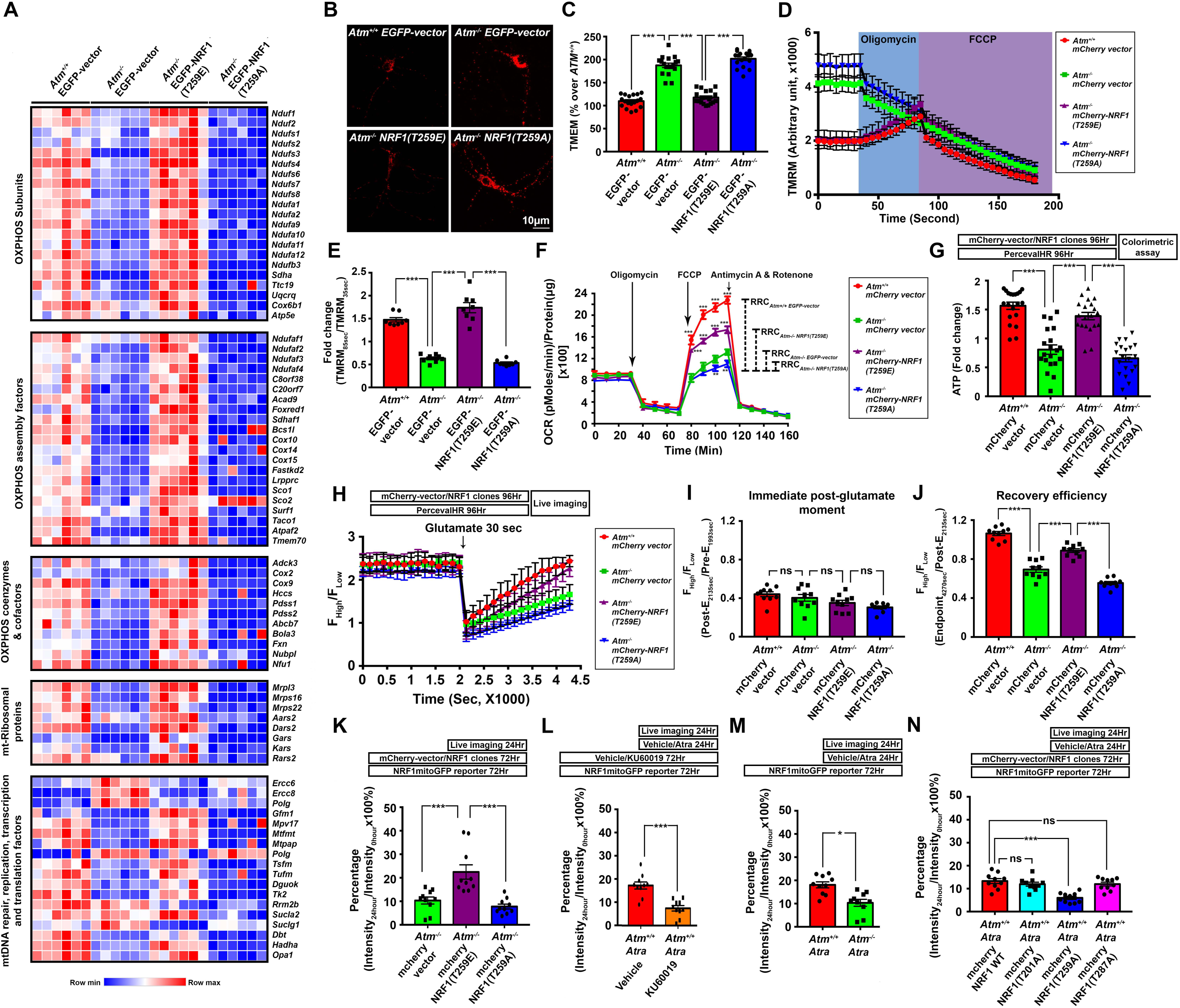
Ectopic expression of T259E NRF1 rescues bioenergetic defects in *Atm*^−/−^ neurons. (A) Heatmaps indicating mRNA expression of 79 nuclear-encoded genes whose loss of function is associated with neurodegeneration. Ectopic expression of T259E but not the T259A mutant of NRF1 for 96 hours significantly rescued the expression of these genes in *Atm*^−/−^ neurons (n=6 biological samples, n=3 technical replicates). (B-C) Static (n=18, *\*\*\*P<0.001*, one-way ANOVA) and (D) dynamic changes in mitochondrial membrane potential in primary cortical neurons were evaluated with TMRM, after overexpressing various forms of NRF1 for 96 hours (n=8). (E) The relative changes in TMRM signals in cells expressing the indicated constructs in the genotypes as shown. TMRM signal was expressed as the ratio of post-oligomycin (85 sec) to pre-oligomycin (35 sec) OCR values (n=8, \*\*\**P<0.001*, one-way ANOVA). (F) Seahorse Mito Stress assay of cellular respiration in neurons pre-expressing the indicated NRF1 constructs for 96 hours (n=10, \*\*\**P<0.001*, *\*\*P<0.01* two-way ANOVA). (G-H) Static and dynamic changes in ATP levels in primary cortical neurons pre-expressing various vectors for 96 hours measured by the Perceval HR reporter. (G) Equilibrium ATP levels at the instant prior the dynamic experiment measured by colorimetric assay (n=20, *\*\*\*P<0.001*, unpaired t-test). (H) Live-imaging of cultured neuron responses to 50 μM glutamate effects on energy consumption. After establishing a steady baseline, glutamate was added for 30 seconds then washed away, and intracellular ATP levels allowed to recover (n=10, *\*\*\*P<0.001*, two-way ANOVA). (I) The relative abundance of ATP:ADP immediately following the removal of glutamate (Signal ratio: Post-E_2135sec_/Pre-E_1993sec_, left panel) and (J) after recovery (Signal ratio: Endpoint_4270sec_/Post-E_2135sec_, right panel) (n=10, *\*\*\*P<0.05*, *ns*=non-significant, unpaired t-test). (K-N) NRF1-mediated transcription activities monitored by the NRF1mitoGFP reporter. (K) *Atm*^−/−^ neurons pre-expressed the indicated forms of NRF1 for 72 hours (n=10, *\*\*\*P<0.001*, one-way ANOVA). Loss of neuronal ATM activity by either (L) treatment of wild type cells with 1 μM KU60019 for 96 hours (n=10, ***P<0.001, unpaired t-test) or (M) treatment of genetically *Atm*^−/−^ cells (n=10, *P<0.05, unpaired t-test) reduced NRF1-reporter activities. (N). Loss of ATM-induced NRF1 reporter activities by 1μM atractyloside treatment (24 hours) is specific to the NRF1-T259A mutant but not point mutants at T201 or T287 (n=10, ns=non-significant, ***P<0.001, one-way ANOVA). For assays in Fig. 9K-M, intracellular GFP signals resulting from NRF1 transcription activity were monitored for 24 hours after the initiation of atractyloside treatments. For Fig.9K-M, ratios of cellular GFP intensities are shown at 24-hour over 0-hour time points.

## Discussion

Our data present an expanded picture of the relationship between the ATM kinase and mitochondrial function and offer a new perspective on the reasons for the particular set of symptoms that are found in persons with A-T. We have found that, separately from the DNA damage response, ATM also functions as a sensor and regulator of a broad cellular response that integrates energy demand and mitochondrial biosynthesis. This newly identified ATM function is particularly important in large, energy intensive cells such as Purkinje cells. The key to the new pathway is the ATM-mediated phosphorylation of NRF1, a transcription factor responsible for the regulation of genes involved in mitochondrial structure and the electron transport chain. A linkage between ATM and mitochondria was first proposed by the Kastan lab (Valentin-Vega et al., 2012) who reported that ATM-deficient fibroblasts had defective mitophagy (the clearance of mitochondria by autophagy), and, as a result, accumulated mitochondria with aberrant morphologies. They identified the Beclin-1 protein, an autophagy inhibitor, as the mediator of the influence of ATM (Valentin-Vega et al., 2012). Fang et al. (2016) extended this line of investigation, showing that defective mitophagy was also found in xeroderma pigmentosa (XPA) cells. They suggested that the problem in XPA lay in PARP1 hyperactivation, which depleted the cellular levels of NAD^+^, and thus reduced the activity of the NAD^+^-responsive histone deacetylase, SIRT1. They proposed that the link between impaired mitophagy and ATM deficiency followed a similar PARP1/NAD^+^/SIRT1 pathway. Other researchers have also reported on the importance of ATM in the regulation of mitochondrial function (Aird et al., 2015; Fang et al., 2016; Morita et al., 2014; Zakikhani et al., 2012). None of these earlier studies, however, focused on the more fundamental linkage between ATM and cellular energy levels.

Our results indicate that a key linkage between ATM and mitochondrial function is through ATP itself, specifically the oxidative stress generated during ATP insufficiency. If TCA cycle components (e.g. acetyl-CoA) decline, mitochondrial NADH_2_ levels drop, the ETC becomes inefficient and the mitochondria cannot sustain an adequate proton gradient across their inner membrane. To protect the proton gradient, which is critical for cellular survival, the ATP synthase transiently runs in reverse and uses ATP (from stores or generated directly from cytosolic metabolism) to pump protons back across the inner mitochondrial membrane, thus reestablishing a healthy Δψm. If this situation persists, non-mitochondrial sources of ATP are insufficient, and a cell responds by increasing its mitochondrial content. This requires enhanced transcription from dozens of nuclear-encoded genes whose protein products collaborate to form the functional electron transport chain. This response is critical to cell viability and thus is regulated by several intersecting pathways which appear to share reactive oxygen species (ROS) as an activating trigger. For example, chronic insufficiency of ATP also results in intracellular calcium increase (Le Masson et al., 2014), which activates multiple extramitochondrial ROS-generating enzymes (Yan et al., 2006). In a similar vein, cytosolic glutathione, a cellular ROS scavenger (Hargreaves et al., 2005), is synthesized *de novo* by two sequential enzymatic ATP-dependent reactions (Chen et al., 2005). In this and possibly other cases, reduced ATP levels lead to increased ROS. It is these intersecting ROS generating pathways that engage ATM in the regulation of cellular metabolism.

We propose that ATM, in its role as a redox sensor is activated by the increased ROS levels caused by the insufficiency of cellular ATP. Activated in this way, ATM phosphorylates the transcription factor NRF1, a master transcriptional regulator of nuclear genes encoding proteins for the mitochondrial respiratory chain (Kelly and Scarpulla, 2004; Scarpulla, 2002). Activated ATM phosphorylates NRF1 at T259 causing the formation of a transcriptionally active NRF1 homodimer. It is highly noteworthy that this pathway of ATM activation is unrelated to its function in the DNA damage response pathway. The downstream targets of the two pathways are different as are the cellular responses. Whether subcellular localization or altered substrate specificity explains these differences remains to be determined. This newly recognized ATM response system is only revealed when the cell is under conditions of stress. While the activities of Complex I-IV are reduced in in the absence of ATM, the affected cells are still able to maintain a nearly normal basal oxygen consumption rate; the steady state leak of protons in *Atm*^−/−^ neurons is also no different from the levels found in wild type cells. It is only when the Δψm is completely depolarized with FCCP or when the cells are challenged with a massive energy demand, such as that following a prolonged depolarization, that the homeostatic failure of *Atm*^−/−^ cells with regard to their ATP levels becomes apparent.

This scenario of sustained ATP deficiency causing ATM-dependent activation of NRF1-related gene transcription offers fresh insight into why cerebellar ataxia, with its regional and cell type specific pathology, manifests as the most prominent neurological phenotype of human A-T. Here the explanation seems to lie in both the physiology of the Purkinje cell and the expression pattern of NRF1. Cerebellar Purkinje cells, with their large cellular size and vast number of synaptic inputs (Carter and Bean, 2009; Sugimori and Llinas, 1990; Welsh et al., 2002), are the only efferent neurons of the cerebellar cortex and are often lost in diseases where ataxia is involved. Computational modeling predicts that each Purkinje cell consumes 62 times more energy than each cerebellar granule cell and 3.9 times more than a cortical pyramidal neuron (Fig.8N, Table S1O). Most of this energy (4.26 × 10^9^ ATP/neuron/s) is used to run the Na^+^/K^+^-pumps that restore the neuronal membrane potential after an action potential (Fig.8N, Table S1O) (Howarth et al., 2012; Howarth et al., 2010). Furthermore, a Purkinje cell typically spikes at high frequency. During a ‘burst’, almost twice as much sodium enters during each spike as would be predicted from the kinetics of a single, isolated action potential (Carter and Bean, 2009). Because of this, a slower-spiking cortical pyramidal neuron uses only 25% more sodium than predicted – closer to the theoretical minimum. This implies that relatively more energy needs to be consumed by the Na^+^/K^+^-pumps to restore ion gradients in Purkinje cells. With their extraordinarily high metabolic/ATP demands, it follows that Purkinje cells are exquisitely vulnerable to oxidative stress and metabolic insufficiency (Sugimori and Llinas, 1990).

This explanation for the unique impact of ATM deficiency on the Purkinje cell is further supported by regional variations in the NRF1 protein itself. Immunocytochemistry has shown that the cerebellum in general and Purkinje cells in particular, have high levels of NRF1 protein. This presumably prepares the Purkinje cell to respond when ATP levels drop, but in the absence of ATM this cellular preparedness cannot be used to increase mitochondrial gene transcription. In keeping with this prediction, a CNS-targeted NRF1 knockout produces Purkinje cell loss, motor ataxia and postnatal lethality (Kobayashi et al., 2011). It is also noteworthy that early onset cerebellar ataxia involving extensive Purkinje cell neurodegeneration is a frequent symptom of patients with mitochondrial DNA defects (Scaglia et al., 2005) or others with mutations of nuclear genes encoding proteins crucial for mitochondrial function (Ashley et al., 2012; Chen et al., 2007). Consistent with our model, these mutations all result in ETC defects (Ashley et al., 2012; Chen et al., 2007; Scaglia et al., 2005).

Our findings can also be seen in the context of the more traditional pathway for the regulation of ATP levels by the AMPK kinase. When AMP levels rise (or ATP levels fall) this heterotrimeric kinase is activated and alters the behavior of a number of downstream targets that enhance catabolic and inhibit anabolic functions, thus allowing the cell to restore ATP levels to normal (recently reviewed in Herzig and Shaw, 2018). Comparing the AMPK response to that of ATM suggests that the former is more likely to respond to moment-to-moment fluctuations in ATP levels, while the latter, with its requirement for significant ROS signaling, ensures a structural and more sustained response to large, chronic changes in ATP. This suggests that ATM functions as a failsafe mechanism that acts as a backup for the AMPK response. When ATP levels suffer a sustained drop to the point where the ETC is compromised and ROS are increased, a cell would be well served by being able to enhance its overall mitochondrial capacity in addition to adjusting the activity of Complex I-V.

Considered together with recent findings on the role of ATM in the modulation of synaptic vesicle trafficking (Cheng et al., 2018), our findings emphasize the importance of the localization, and independent functions, of ATM in cytoplasm and nucleus. Although it has been suggested that the neurodegenerative phenotype of A-T is mainly attributable to inefficient DNA repair, it has always been challenging to explain the full range of neurological and behavioral symptoms by invoking only this pathway. A-T cells show normal levels of DNA repair (Kastan and Lim, 2000) although the process is slower. Further, since DNA damage is mostly a stochastic event that occurs throughout the genome of all cells, it is difficult to explain how inefficient repair of genome-wide DNA damage explains the selective loss of Purkinje cells in all A-T patients.

Combining the predicted energy demands unique to Purkinje cells with a defective capacity to respond to the drop in ATP levels brought on by high levels of neuronal activity, makes the neurological symptoms of A-T seem much less puzzling.

## Materials and methods

### Mitochondrial database

Associations between various symptoms of ataxia-telangiectasia were generated using the online database http://www.mitodb.com (Scheibye-Knudsen et al., 2013). On the home page of the database, ataxia-telangiectasia was chosen as the disease to be tested. Once test results were generated, the number of mitochondrial versus non-mitochondrial diseases (including those in the Unknown category) associated with different symptoms of ataxia-telangiectasia were tabulated on the subpage named “Barcode”. Details of the data can be found in Table S1A.

### Multiple protein sequence alignment

Protein sequences of NRF1 in *Homo sapiens* (human) [NP_001035199.1], *Pan troglodytes* (chimpanzee) [XP_001155812], *Mus musculus* (mouse) [NP_035068.3] and *Rattus norvegicus* (rat) [NP_001094178.1] were obtained from the NCBIGene database https://www.ncbi.nlm.nih.gov/gene. Multiple protein sequence alignment was performed on the Clustal Omega platform https://www.ebi.ac.uk/Tools/msa/clustalo. Conserved (S/T)Q sites among all the species analyzed were manually identified for subsequent analysis.

### Human microarray data mining

A human expression array dataset (GEO Accession GSE61019), comparing samples of A-T and control cerebella (Jiang et al., 2015), was analyzed with GEO2R. Raw data from each of the samples were extracted.

#### Gene Ontology (GO) analysis

All transcripts that reflected significant changes (Adjusted p-value  0.05) by the GEO2R analyzer were extracted (Table S1B, yellow highlights) and analyzed on the GATHER platform http://changlab.uth.tmc.edu/gather/gather.py (Chang and Nevins, 2006). Among all 30 GO terms that reflected the greatest possible significance (p<.0001), we performed enrichment analysis (listed in descending order) based on the corresponding Bayes factors and the number of genes indicated in each GO term.

#### Gene Set Enrichment analysis (GSEA)

We performed a second analysis using GSEA with the KEGG pathway groupings of genes http://software.broadinstitute.org/gsea/index.jsp (Subramanian et al., 2005). The full list of phenotypes enriched in the normal can be found in Table S1E. Pathways with normalized p-value (NOM p-val) < 0.01 are highlighted in yellow.

#### Human MitoCarta2.0 analysis

We performed a third analysis using MitoCarta2.0 *—* a collection of 1,158 nuclear and mitochondrial genes encoding proteins with strong evidence for mitochondrial localization https://www.broadinstitute.org/files/shared/metabolism/mitocarta/human.mitocarta2.0.html (Calvo et al., 2016). The matching genes were extracted from the raw dataset of this human array for expression analysis. The full list of extracted data from each individual sample can be found in Table S1F. Table S1G contains the comparison data between the control versus A-T groups. Specific genome localization of each gene was also determined. Note that a list of 42 genes from the MitoCarta2.0 list failed to match probes on the human expression array.

#### PSCAN gene promoter analysis

To identify potential transcription factors that result in global changes of gene expression, we used the PSCAN platform http://159.149.160.88/pscan/ (Zambelli et al., 2009) to analyze the genes identified in the human expression array data that showed significant differences of expression between A-T and control cells. The list of gene IDs was first converted to the corresponding RefSeq mRNA IDs using the DAVID platform https://david.ncifcrf.gov (Huang da et al., 2009), and subsequently input into PSCAN using the “human and mouse” species option; a default promoter setting of “-450 to +50” was used to cover the regions located immediately adjacent to the transcription start site. The JASPAR 2014 matrices were also selected for analysis. The results are compiled in Supplementary Table S2. Detailed matrix information, sample mean score and statistics are listed for each site of interest. NRF1 was the gene that ranked highest in this analysis and was thus subjected to further scrutiny. The list of NRF1 binding sites in each input promoter, with corresponding gene name, oligo score, position with respect to transcription start site of the gene, oligo sequence and strand were extracted for further analysis.

### *In silico* protein structure modeling, docking, and contact area analysis

With the protein sequence of the human NRF1 protein [NP_001035199.1] obtained from the NCBI-Gene database, the DNA binding and dimerization domain (aa 116-283) was used to predict the corresponding 3- D protein structure on the Phyre^2^ engine http://www.sbg.bio.ic.ac.uk/∼phyre2/html/page.cgi?id=index using the intensive modelling mode (Kelley et al., 2015). The highest-ranking model (with score > 85), was used to overlap the excised bromodomain structures of BRD2 (PDB: 5UEW) and BRD4 (PDB: 4O76) on Molsoft ICM-Pro Browser, or directly adopted for subsequent *in silico* docking analysis for the homodimerized NRF1 structure on the ZDOCK server http://zdock.umassmed.edu (Pierce et al., 2014). The best-ranked model (with the highest ZDOCK score) from each analysis was then subjected to contact area analysis for visualizing interacting amino acid residues with the Molsoft ICM-Pro Browser. To evaluate the monomer structures, T259 was mutated to T259(PO_4_^−^) using Pymol mutagenesis tool prior to the same docking procedures and contact area analysis.

### Animal maintenance, brain tissue isolation, primary neuronal and mouse embryonic fibroblast (MEF) culture

C57BL/6J and 129S6/SvEvTac-Atmtm1bal/J (heterozygous for an engineered ATM kinase domain mutation – Xu et al., 1996) were obtained from The Jackson Laboratory. Colonies were maintained in the Animal and Plant Care Facility of the Hong Kong University of Science and Technology (HKUST). Homozygous mutants (*Atm*^−/−^) were generated by mating heterozygous carriers (*Atm*^+/−^), as per The Jackson Laboratory protocol. All animal experimental protocols were approved by the Animal Ethics Committee at HKUST and their care was in accord with the institutional and Hong Kong guidelines

Brain tissue was isolated by anesthetizing adult mice with intraperitoneal administration of 1.25% (vol/vol) Avertin at a dosage of 30 mL/kg body weight. The heart of each mouse was then surgically exposed, the left chamber was catheterized, and the right atrium was opened. Chilled physiological saline was perfused transcardially for 3 minutes to remove blood from the body. After perfusion, the cranial bones were opened; cortex and cerebellum were harvested, snap-frozen in liquid nitrogen, and stored at 80 °C until use.

Embryonic cortical neurons were isolated by standard procedures (Chow et al., 2014). Gravid females were killed on the 16^th^ day of gestation and the E16.5 embryos collected in ice-cold PBS‐ glucose. The cortical lobes were removed, following which the meninges were removed and the cortices were placed in 1X trypsin solution for 10 minutes, with manual shaking at 5 min. After digestion, an equal volume of DMEM with 10% (vol/vol) FBS was added to inactivate the trypsin. Samples were then centrifuged at 2,500 rpm for 5 min. Supernatant was removed, followed by transferring the pellet to fresh Neurobasal medium supplemented with B-27, penicillin/streptomycin (1X) and L-glutamine (2mM; GlutaMAX, Invitrogen) prior to gentle resuspension. Tissue was triturated 10 times through a 5-mL pipette and allowed to settle to the bottom of a 15-mL conical tube. Cells in solution above the pellet were removed. Surviving cells were identified by trypan blue exclusion and counted before plating on poly-L-lysine-coated (0.05 mg/mL) glass coverslips. Unless otherwise specified, cells were plated in 24-well plates at 50,000 cells per well and allowed to mature for over 7-10 days in vitro (DIV) before transfection or lentiviral transduction. For other experiments, cells were grown for a minimum of 14 days in vitro (DIV14) before drug-treatment experiments. Every 3 days half of the culture medium was replaced with an equal volume of fresh medium to sustain the culture.

MEF cell cultures were established from the same E16 embryos used for isolation and culture of primary cortical neurons. After the heads (for primary neuronal culture), tails, limbs, and internal organs were removed, the remaining tissue was minced and trypsinized for 20 min, then seeded into P100 culture dishes in 12 mL of the complete MEF medium [Iscove’s modified Dulbecco medium (IMDM) containing 10% iron-supplied calf serum (Hyclone), with additional nonessential amino acids (NEAA), 2 mM GlutaMAX^TM^ and 1% Penicillin-Streptomycin.]. When cultures became confluent, the cells were split 1:2 or 1:3 then passaged two or three times to obtain a morphologically homogenous culture. At this point cells were harvested, and separated into aliquots that were either frozen or expanded for further studies.

### Postmortem human brain samples

Frozen postmortem brain samples of diseased A-T patients and age-matched controls were requested from the NeuroBioBank, NIH. All samples were thoroughly anonymized prior distribution. The US Human Studies Board characterizes these tissues as Exempt (EXMT). All studies conducted using human tissue were strictly complied with the ethical standard of NIH and the Ethics Committee on Research Practices of the Hong Kong University of Science and Technology. Frozen sections were prepared from brain blocks and were used for immunohistochemistry purposes.

### Cell culture and transfection

Lines of primary human skin fibroblasts from A-T patients and their respective controls were obtained from the NIA Aging Cell Repository of the Coriell Institute for Medical Research and maintained in DMEM/10% FBS. Primary neurons were isolated from E16 embryos of C57BL/6J or *Atm*^+/−^ pregnant mice and cultured as described above. DNA constructs were transfected with Lipofectamine LTX in the presence of Plus Reagent into both types of primary culture following the manufacturer’s protocol. At 5 hours after transfection, cells were refreshed with new culture medium (or conditioned medium for primary neurons) and further incubated for 48-72 hours to allow recovery and ectopic expression of transfected constructs.

### Lentivirus production and transduction

Lentivirus stocks were produced as previously described with slight modifications (Cribbs et al., 2013). Human embryonic kidney 293FT cells (Invitrogen) were transfected using Lipofectamine 2000 (Invitrogen) with the expression of two helper plasmids: psPAX2 and pMD2.G. Ten micrograms of the transfer vector, 5 μg pMD2.G and 5 μg psPAX2 of DNA were used per 10-cm plate. Forty-eight hours after transfection, the supernatants of four plates were pooled, centrifuged at 780 × g for 5 minutes, filtered through a 0.45-μm pore size filter, and further centrifuged at 24,000 rpm for 2 hours. The resulting pellet was resuspended in 100 μL of PBS. Lentivirus titration was performed with a p24 ELISA (Clontech).

### Co-immunoprecipitation, SDS-PAGE-Western Blotting and native PAGE-Western blotting

Isolated brain tissues or cell pellets were homogenized in RIPA buffer (Millipore) with 1X complete protease inhibitor mixture (Roche) and 1X PhosSTOP phosphatase inhibitor mixture (Roche) on ice then centrifuged for 10 minutes at 18,400 × g to remove large debris. The protein concentration of the supernatant was determined by Bradford Assay (Bio-Rad). For coimmunoprecipitation, 1 mg of the total cell lysates was first incubated with control IgG (Santa Cruz Biotechnology) for 30 minutes, pre-cleared with 50 μL Dynabeads Protein G (Invitrogen), and then incubated with various antibodies overnight at 4 °C, using the suggested dilutions from the product datasheets. Beads bound with immune complexes were collected by DynaMag-2 (Life Technologies) and washed three times before elution in 90 μL of buffer containing 0.2M Glycine-HCl, pH 2.5, which was neutralized with 10 μL of neutralization buffer [1M Tris-HCl (pH 9.0)]. The eluates were subjected to 9-15% SDS-PAGE and Western blot analysis.

For SDS-PAGE (polyacrylamide gel electrophoresis), 100 μg of proteins derived from cell or tissue lysates were prepared in 5X sample buffer [10%w/v SDS; 10mM beta-mercaptoethanol; 20%v/v glycerol; 0.2M Tris-HCl, pH6.8; 0.05% Bromophenol blue]. With a Bio-Rad system, separating gels of different acrylamide percentages (6%-15%) were prepared with the following components in double distilled water: acrylamide/bis-acrylamide (30%/0.8% w/v); 1.5M Tris (pH=8.8); 10% (w/v) SDS; 10% (w/v) ammonium persulfate and TEMED) and a 5% stacking gel (5 ml prep: 2.975ml Water; 1.25 ml 0.5M Tris-HCl, pH6.8; 0.05 ml 10% (w/v) SDS; 0.67ml acrylamide/bis-acrylamide (30%/0.8% w/v); 0.05ml 10%(w/v) ammonium persulfate and 0.005 ml TEMED) were prepared. Samples were run in SDS-containing running buffer [25mM Tris-HCl; 200mM Glycine and 0.1% (w/v) SDS] until the dye front and the protein marker reached the foot of the glass plate. Standard immune-blotting procedures were used, which include protein transfer to polyvinylidene difluoride (PVDF) membranes, blocking with non-fat milk, and incubation with primary and secondary antibody, followed by visualization with the SuperSignal West Dura/Femto Chemiluminescent Substrate (ThermoFisher Scientific).

For native-PAGE, the same discontinuous chloride and glycine ion fronts as SDS-PAGE were used to form moving boundaries that stack and then separate polypeptides by their charge to mass ratio. Proteins (100 μg) were prepared in a non-reducing non-denaturing sample buffer (2X: 62.5 mM Tris-HCl, pH 6.8; 25% glycerol; 1% bromophenol blue) without heating to maintain secondary structure and native charge density. With a Bio-Rad system, 10 ml/gel 8% native PAGE separating gel [2.6ml acrylamide/bis-acrylamide (30%/0.8% w/v); 7.29ml 1.5M Tris-HCl (pH8.8); 100 μl 10%(w/v) ammonium persulfate and 10 μl TEMED) and 5 ml/gel native PAGE stacking gel [4.275 ml 0.5M Tris-HCl. pH=6.8; 0.67ml acrylamide/bis-acrylamide (30%/0.8% w/v); 0.05ml 10%(w/v) ammonium persulfate; 5μl TEMED] were prepared. Samples were run in a non-denaturing running buffer (25mM Tris, 192 mM glycine, pH 8.3) with the entire system cooled with ice. After electrophoresis, standard immune-blotting procedures were followed as above.

### Quantitative RT-PCR (qPCR) analysis

Total cellular RNA was purified from brain cortex tissues or cultured cells using the RNeasy mini kit (Qiagen) following the manufacturer’s protocol. For quantitative real-time PCR (qPCR), RNA was reverse-transcribed using the High-Capacity cDNA Reverse Transcription Kit (Applied Biosystems) according to the manufacturer’s instructions. The resulting cDNA was analyzed by qRT-PCR using SYBR Green PCR Master Mix (Applied Biosystems). All reactions were performed in a Roche LightCycler (LC) 480 instrument using the following protocol: pre-incubation at 95 °C for 15 min (1 cycle); denaturation at 94 °C for 15 seconds, annealing and extension at 55 °C for 30 seconds (40 cycles), melting at 95 °C for 5 seconds, 65 °C for 60 seconds and 95 °C continues (1 cycle) followed by cooling at 40 °C for 30 seconds. The specificity of the primers was confirmed by observing a single melting point peak. qPCR efficiency was calculated from the slope was between 95 and 105% with co-efficiency of reaction R^2^=0.98-0.99. A total of 7-9 biological replicates × 4 technical replicates were performed for each treatment group. Data was analyzed using the comparative Ct method (ΔΔCt method).

### Mitochondrial complex activity assays

Activities of Complexes I to IV were measured using kits purchased from BioVision (K968-100, K660-100, K520-100 and K287-100) and that of Complex V was measured using a kit from Invitrogen (KHM2051). Measurements were conducted by strictly adhering to the manufacturers’ instructions.

### Imaging intracellular pyruvate level in single cells with the FRET sensors, Pyronic and Laconic

Pyruvate sensor, Pyronic (Pyruvate Optical Nano-Indicator from CECs), was obtained from Addgene (Plasmid #51308) as was the lactate sensor, Laconic (LACtate Optical Nano Indicator from CECs) was obtained from Addgene (Plasmid #44238). For both sensors, the exposure of the sensor to its ligand (pyruvate or lactate) causes an increase in mTFP (donor) fluorescence intensity and a decrease in Venus (mCFP, receiver) fluorescence intensity, thus resulting in a decrease in FRET efficiency. As reported previously (San Martin et al., 2014), the biosensors were transfected into cortical neurons on DIV9. Twenty-four hours after transfection, cells were exposed to one of the following drug conditions:

> *For Pyronic:*
>
> i) vehicle (saline); ii) 30 mM 2-deoxyglucose (2-DG) (Saraiva et al., 2010); iii) 1 μM UK5099 (Jourdain et al., 2016) or iv) 2.5mM oxamate (Choi et al., 2012)] in KRH-bicarbonate buffer (112mM NaCl, 1.25mM CaCl_2_, 1.25mM MgSO_4_, 1-2 mM glucose, 10mM HEPES, 24mM NaHCO_3_ (pH7.4) and 3mM KCl).
>
> *For Laconic:*
>
> i) vehicle (saline); ii) 500 μM pCMBS (San Martin et al., 2014); or iii) 2.5 mM Oxamate (Choi et al., 2012)] in imaging buffer (112mM NaCl, 1.25mM CaCl2, 1.25mM MgSO4, 1–2mM glucose, 10mM HEPES, 24mM NaHCO3, pH 7.4, with 3 mM KCl).

Cultures were then live-imaged for another 24 hours with imaging at 6-hour intervals in a 95% air/5% CO2-gassed incubator using a Leica SP8 confocal microscope equipped with a 63X oil-immersion objective. Image acquisition was controlled by the Leica Application Suite X Microscope Software (LAS X). To perform nanosensor ratio measurements, cells were excited at 430 nm for 0.2-0.8 seconds. Emission was divided with an Acousto-Optical Beam Splitter (Leica Microsystems Inc.), and recorded at 480±20 (mTFP) and 535±15nm (Venus). The ratio between mTFP and Venus along time was used to estimate pyruvate concentrations inside the cell.

### Colorimetric assay

Cellular contents of pyruvate were measured using kits purchased from BioVision (K609-100) following the manufacturers’ instructions. Cellular contents of lactate were measured using specific kits purchased from BioVision (K607-100), following the manufacturer’s instructions.

### Cytosol and mitochondria separation followed by isocitrate and succinate colorimetric assays

pMX-3XFLAG-EGFP-OMP25 plasmid was obtained from Addgene (Plasmid #83354). Human primary fibroblasts were first transfected with the plasmid as described above. In cases where the transfection efficiencies were 80% or greater, after 72 hours mitochondria were harvested by immunopurification (Chen et al., 2016b). 3X-FLAG-EGFP-OMP25-expressing cells were gently scraped into 1 ml KPBS (136 mM KCl, 10 mM KH_2_PO_4_, pH 7.25). The cell suspension was centrifuged at 1000 × g for 2 minutes, supernatant was discarded, and the cell pellet gently resuspended in 1 ml of the kit-supplied assay buffer. Cells were homogenized with 25 strokes in a 2 ml Dounce homogenizer, taking care not to introduce air bubbles into the solution. The homogenate was then centrifuged and the mitochondria isolated by anti-FLAG IP (the entire procedure was performed on ice in a 4°C cold room with pre-chilled buffers and equipment). To increase the speed of isolation only one sample for one metabolite analysis was processed at a time. Anti-FLAG magnetic beads (200 μl) were pre-washed three times with KPBS by gentle pipetting using a wide-bore pipette tip. Beads were collected using a DynaMag (Thermo Fisher Scientific). Sample was then added to the beads and the mixture incubated overnight. The following day antibody-bounded mitochondria were separated from the cytosol using the DynaMag; the cytosol fraction was saved for metabolite analysis. The isolated beads were quickly washed twice with 1 ml KPBS, re-suspended in 1 ml assay buffer, and transferred to a new tube. An aliquot of the mitochondrial suspension (250 μl) was taken for detergent lysis and protein quantification; the remainder was used for metabolite extraction analyses.

Cytosolic and mitochondrial contents of isocitrate and succinate were measured using kits purchased from BioVision (K656-100 and K649-100 respectively).

### Live imaging of cytoplasmic redox status

Cyto-roGFP obtained from Addgene (Plasmid #49435) senses redox changes in a cell (Waypa et al., 2010). The Cyto-roGFP biosensor was transfected into cortical neurons on DIV9, and 24 hours after transfection, neurons were exposed to one of the following conditions: i) vehicle (saline); ii) 10 nM H_2_O_2_ (Wang et al., 2007); or iii) 1 μM atractyloside (Schutt et al., 2012) in Neurobasal culture medium. Cultures were then imaged for 24 hours at hourly intervals in a 95% air/5% CO2-gassed incubator using a Leica SP8 confocal microscope equipped with a 63X oil-immersion objective, controlled by the Leica Application Suite X Microscope Software (LAS X). The redox-sensitive protein reporter has excitation maxima at 400±15nm and 484±15nm, and an emission maximum at 525±15nm. The relative amplitudes of these peaks depend on the state of oxidation. With increased oxidation the 400±15nm excitation peak increases while the 484±15nm peak decreases (Hanson et al., 2004), Importantly, the ratiometric nature of the analysis renders the results independent of the expression levels of the plasmid in any one cell. Data were collected with the Leica Application Suite X Microscope Software. The fluorescence excitation ratios were obtained by dividing integrated intensities obtained from manually selected portions of the imaged regions of intact whole cells collected using 400±15nm and 480±15nm excitation filters after appropriate background correction. Background correction was performed by subtracting the intensity of a nearby cell-free region from the signal of the imaged cell.

### Imaging intracellular ATP-to-ADP ratio

Perceval High Range (PercevalHR) obtained from Addgene (Plasmid #49082) senses changes in the ATP:ADP ratios in mammalian cells (Tantama et al., 2013). We transfected PercevalHR into cortical neurons on DIV9. Twenty-four hours after transfection, neurons were imaged on a Carl Zeiss AxioOberver Z1 Inverted Microscope with a piezo Z stage glass slide insert. The sensor has two distinct peaks of excitation, ATP binding increases the fluorescence at 500 nm, while ADP increases fluorescence at 420 nm. PercevalHR was excited sequentially using 482±15nm and 445±15nm band pass filters; emission was measured with a 529±15nm band pass filter. The excitation and emission signals were optically separated with a 490 nm short pass dichroic mirror. Images were taken over a period of 72 minutes (capture time, 2 min). The ratio of the fluorescence signal at the two different excitation wavelengths (F_high_/F_low_), reports the ATP occupancy of the reporter, and is independent of the amount of sensor protein in the cell.

During the imaging process, neurons were placed in a custom chamber which was continuously perfused with a bathing solution (2 ml/min; 32-34°C) containing 120mM NaCl, 3mM KCl, 2mM CaCl_2_, 1mM MgCl_2_, 3mM NaHCO_3_, 1.25mM NaH2PO4, 15mM HEPES, 5mM glucose, 0.2mM pyruvate, and 0.5mM GlutaMax adjusted to pH 7.4. We determined a baseline reporter signal by imaging a resting cell for at least 25 minutes. After ensuring a stable signal, 50 μM glutamate was applied for 30 seconds to strongly excite the neurons. Following the depolarization, the cell would need considerable amounts of ATP to drive the Na^+^/K^+^-ATPase (sodium pump) to be able to restore the resting ion gradients. We followed the drop in ATP and its subsequent restoration by the mitochondria for 30-45 minutes after the glutamate stimulus.

### ATP colorimetric assay

The cellular contents of ATP were measured using kits purchased from BioVision (K354-100).

### Determination of mitochondrial membrane potential

Tetramethylrhodamine, methyl ester (TMRM, Thermofisher) is a cell-permeant, cationic, red-orange fluorescent dye that is sensitive to the mitochondrial membrane potential; Mitotracker green (Thermofisher) – a mitochondrial dye that is insensitive to Δψm – was used as a counterstain. Stock solutions of the dyes (10mM) were prepared in anhydrous dimethylsulfoxide (DMSO). Cultured cortical neurons or human fibroblasts were first washed thrice with Tyrode’s buffer (145 mM NaCl, 5 mM KCl, 10 mM glucose, 1.5 mM CaCl_2_, 1 mM MgCl_2_, and 10 mM HEPES; adjusted to pH 7.4 with NaOH). Working concentrations (20 nM) of TMRM and Mitotracker green were prepared in Tyrode’s buffer and applied to the cells. After incubation in dark at room temperature for 45 minutes, imaging was performed on a confocal microscope (Leica, SP8) using the time-series and sequential scanning programs. TMRM was excited at 514±15nm and the emission at 570±15nm was measured. Mitotracker green was excited at 480±15nm and detected at 520±15nm. To test changes in mitochondrial membrane potential, 1 μM carbonyl cyanide 4-(trifluoromethoxy)phenylhydrazone (FCCP) or 2 μg/ml of oligomycin, were applied, which depolarized or hyperpolarized the mitochondrial membrane potential, respectively (Joshi and Bakowska, 2011).

### Assessment of mitochondrial function

The mitochondrial oxygen consumption rate (OCR) in both primary neurons and human fibroblasts were assessed using a Seahorse Bioscience XF^e^24 analyzer (Agilent, Massachusetts, USA) in combination with the Seahorse Bioscience XF Cell Mito Stress Test assay kit. In this assay, sequential additions of the ATP synthase inhibitor, oligomycin, the mitochondrial uncoupler, carbonyl cyanide 4-(trifluoromethoxy)-phenylhydrazone (FCCP) and the complex I + II inhibitors rotenone + antimycin A provide insight into different aspects of mitochondrial function as described in the text. Experiments were performed on intact adherent cells.

#### Coating and cell plating

The surface area of each well of the 24-well Seahorse XF24 microplate is identical to that of standard 96-well microplates (0.32 cm^2^). Poly-L-lysine (0.1 mg/ml in sterile borate buffer [pH8.4], 500 μl/well) was incubated overnight at 37°C, 5% CO_2_, and 95% humidity. The substrate solution was then removed, and the microplates were dried without further rinsing under a sterile laminar flow hood. Dissociated primary cortical neurons were isolated as described above. 100,000 neurons/well or 80,000 primary fibroblasts/well were seeded in a volume of 150 μl. Immediate after seeding, plates were incubated for 1 hour at 37 °C prior bringing the total volume of each well to 500 μl with corresponding culturing medium (Neurobasal medium for neurons; DMEM with 10% FBS for human fibroblast). Fibroblast cultures were maintained for 2 days prior to analysis. For neurons, after 1 night of settling, cultures were treated with 20 μM FDU in Neurobasal medium for 24 hours with medium changed every other day; this method eliminates most of the non-neuronal cells. Neurons were maintained until DIV7 prior to subsequent analysis.

#### Cell mito stress test assay

Growth medium was removed, leaving 50 μl in each well after which cells were rinsed twice with 400 μl of pre-warmed assay medium consisting of XF base medium supplemented with 25 mM glucose, 2 mM glutamine and 1 mM sodium pyruvate buffered to pH7.4. Following the rinses, 475 μl assay medium and 50 μl conditioned medium (525 μl final) were added to each well. Cells were then incubated in a 37 ^°^C incubator without CO_2_ for 1 hour after which pre-warmed oligomycin, FCCP, rotenone and antimycin A solution were loaded into injector ports A, B and C of the sensor cartridge to achieve final concentrations of 0.5 μM for oligomycin, 1 μM for FCCP and 1 μM for rotenone and antimycin A. The cartridge was calibrated with the XF24 analyzer, and the assay performed as described (Nicholls et al., 2010).

The oxygen consumption rate (OCR) was measured and changes from baseline rates were tracked following the sequential addition of oligomycin, FCCP and the rotenone plus antimycin A mixture. This allowed for calculation of the basal respiration rate, proton leakage, maximal respiration, spare respiratory capacity, non-mitochondrial respiration and ATP production (Nicholls et al., 2010) by Seahorse XF24 software version 1.8. At the end of each assay, cells were washed once with an excess of room temperature PBS, lysed with ice-cold RIPA buffer (0.15 M NaCl, 1 mM EDTA, 1 mM EGTA, 0.5% sodium deoxycholate, 0.1% SDS, 1% Triton X-100, 50 mM Tris-HCl [pH 8] with added protease and phosphatase inhibitor cocktails). Protein content was estimated by the Bio-Rad Dc protein assay (Bio-Rad, Hercules, CA, USA) using a Molecular Devices Softmax M3 microplate reader (Sunnyvale, CA, USA). Data from each well were then normalized for total protein content. Normalized Seahorse XFe24 measurements were used to calculate a mean (± SEM) for each density group and each plate using Seahorse XF24 software version 1.8 (Seahorse Bioscience, Billerica, MA, USA).

#### Metabolic Fuel Flux Assays

The Mito Fuel Flex Tests were performed on Seahorse XFe24 Bioanalyzer (Agilent). All assays were performed following manufacture’s protocols. Briefly, the test inhibits import of three major metabolic substrates (pyruvate, fatty acids, and/or glutamine) with mitochondrial pyruvate carrier inhibitor UK5099 (2 μM), carnitine palmitoyltransferase 1A inhibitor etomoxir (4 μM), or glutaminase inhibitor BPTES (3 μM). This test determines cellular dependence on each of the metabolites to fuel mitochondrial metabolism by inhibiting the individual substrate import. Baseline OCR was monitored for 18 minutes followed by sequential inhibitor injections (i.e. Treatment 1 or Treatment 2) with OCR reading for 1 hour following each treatment. The inhibitor treatments and calculations are shown below:

Dependency (%)=[(Baseline OCR-Target inhibitor OCR)/(Baseline OCR-All inhibitors OCR)]x100%

| Metabolic tests and associated inhibitors |  |  |
| --- | --- | --- |
| Metabolite test | Treatment 1 | Treatment 2 |
| Glucose/Pyruvate dependence | UK5099 | Etomoxir + BPTES |
| Glutamine dependence | BPTES | Etomoxir + UK5099 |
| Fatty acid dependence | Etomoxir | BPTES + UK5099 |

#### Immunocytochemistry

For immunostaining, primary neuronal cultures were grown on 13-mm coverslips (Marienfeld) in 24-well plates. At the appropriate times, cultures were fixed with fresh 4% (wt/vol) paraformaldehyde (Sigma-Aldrich) for 10 min, and the cells permeabilized by treatment with 0.3% Triton X-100 in PBS for 10 min. After blocking with 5% (wt/vol) BSA in PBS for 1 hour, primary antibodies were added and incubated overnight at 4 °C. The following day, coverslips were washed three times (10 min each) with PBS. After rinsing, secondary antibodies were applied for 1 h at room temperature followed by three additional washes with PBS. The coverslips were then inverted and mounted on glass slides with ProLong Gold Antifade Reagent (Life Technologies). Immunofluorescence was analyzed and Z-stack maximum projected images were photographed using a TCS SP8 confocal microscope (Leica Microsystems Inc.).

For immunostaining of brain sections, we performed antigen retrieval by treating the mounted cryostat sections with citrate buffer at 95 °C for 15 min followed by cooling to room temperature for 1 hour. Sections were rinsed in PBS, blocked with 5% (vol/vol) donkey serum/PBS for 1 hour before immunostaining as above.

#### LIVE/DEAD cell viability assays

The LIVE/DEAD viability/cytotoxicity kit for mammalian cells was purchased from Thermofisher. The method relies on two probes — calcein AM and ethidium homodimer-1 (EthD-1). Cultured cortical neurons or human fibroblasts were washed three times with 500 volumes of Dulbecco’s phosphate-buffered saline (D-PBS) to remove or dilute serum esterase activity generally present in serum-supplemented growth media. Two hundred microliters of the calcein/EthD-1 working solution (1 μM and 2 μM respectively in D-PBS) were applied to the surface of a coverslip or 35-mm disposable petri-dish and incubated for 45 minutes at room temperature in the dark. Imaging was performed on a confocal laser microscope (Leica, SP8), with the sequential scanning program. Calcein was excited at 485±15nm and emission was measured at 530±15 nm; EthD-1 was excited at 530±15nm with emission measured at 645±15nm. Quantification of live (green) and dead (red) cells was performed randomly on 10 fields from each of 3-4 blinded samples. The percentage of dead cells was calculated as (number of dead cells/total number of live and dead cells) × 100%.

#### Monitoring NRF1 nuclear transcription activity and changes in mitochondrial morphology

The NRF1mitoGFP reporter was a kind gift from Prof. Johan Tronstad (Hodneland Nilsson et al., 2015). Cortical neurons were transfected with reporter-expressing plasmids on DIV9. Twenty-four hours after transfection, cells were exposed to one of the following drug conditions i) vehicle (saline); ii) 1 μM atractyloside alone or iii) atractyloside plus 5 mM N-acetylcysteine (NAC) (Chao et al., 2016) followed by live-imaging for another 24 hours in a 95% air/5% CO_2_-gassed chamber mounted on a Leica SP8 confocal microscope equipped with a 63X oil-immersion objective, controlled by the Leica Application Suite X Microscope Software (LAS X). GFP was excited at 480±15nm and emission detected at 520±15nm. To analyze mitochondrial morphology the GFP signals of each cell (in a different set of cultures) were imaged at 24 hours after drug treatment and later the Z-stack images were used to construct the 3D model by the Leica Application Suite X Microscope Software (LAS X) for morphology analysis.

#### Nrf1 expression plasmid and mutated Nrf1

A mouse cDNA ORF clone encoding NM_001164226, tagged with Myc-FLAG at the C-terminus, was purchased from OriGene (catalog number MR226726). EGFP or mCherry coding sequences were fused to the C-terminus, replacing the Myc-FLAG tag, by Gibson assembly to make pCMV6-Nrf1-EGFP. Site-directed mutagenesis used the NEBaseChanger kit (NEB), with oligonucleotide primers designed using the manufacturer’s website. All mutations and deletions were confirmed by sequencing.

#### Quantification of the relative abundance of mouse mitochondrial DNA to nuclear DNA

The quantification protocol for mouse mitochondrial DNA without co-amplification of nuclear mitochondrial insertion sequences was adopted (Malik et al., 2016). To prepare genomic DNA, mouse tissue samples were homogenized using TissueLyser (Qiagen) to eliminate cross-contamination. Total genomic DNA was extracted using the DNA easy Blood & Tissue kit according to the manufacturer’s instruction (Qiagen). Before proceeding to qPCR, the DNA template was subjected to pre-treatment shearing (Bath sonicator Kerry, Pulsatron 55 – 38kHz for 10 minutes) as described previously in order to avoid dilution bias (Malik et al., 2011). The template concentration was determined using NanoDrop and adjusted to 10 ng/μl. To avoid errors arising from repeated freeze thaw cycles DNA samples were kept at 4°C for the duration of study.

The mitochondrial DNA content was assessed by absolute quantification using real time PCR. Primers of mouse mitochondrial DNA (mMito) and mouse single copy nuclear β-2 microglobulin (B2M) were used to amplify the respective products. PCR products were purified and used to prepare dilution standards for both amplicons using a range of dilutions from 10^2^−10^8^ copies per 2 μl for quantification. Mitochondrial copy number per cell was determined by carrying out qPCR in a total volume of 10 μl, containing 5 μl of Quantifast SYBR Master Mix (Qiagen), 0.5 μl of forward and reverse primer (400 nM final concentration each), 2 μl template DNA and 2 μl DNase free water. The reactions were performed in a Roche LightCycler (LC) 480 instrument using the following protocol: pre-incubation at 95 °C for 5 min (1 cycle); denaturation at 95 °C for 10 seconds, annealing and extension at 60 °C for 30 seconds (40 cycles), melting at 95 °C for 5 seconds, 65 °C for 60 seconds and 95 °C continues (1 cycle), cooling at 40 °C for 30 seconds. The specificity of the primers was confirmed by the presence of a single melt peak. qPCR efficiency calculated from the slope between 95 and 105% with co-efficient of reaction, R^2^=0.98-0.99. A total of 7-9 biological replicates × 4 technical replicates were performed for each treatment group. Data was analyzed using the comparative Ct method (ΔΔCt method).

## Reagents and resources

### Antibodies

Anti-Actin (Cell Signaling Technology, #4970)

Anti-ATM (phospho S1981)

(Abcam, ab81292)

Anti-FLAG M2 (Sigma, F3165)

Anti-Gamma-H2AX (phospho S139) (Abcam, ab2893)

Anti-GFP (Santa Cruz Biotechnology, sc-8334)

Anti-MAP2 (Abcam, ab5392)

Anti-Myc-Tag (Cell Signaling Technology, #2276)

Anti-NRF1 (ThermoFisher Scientific, PA5-40912)

Anti-Phospho-(Ser/Thr)

ATM/ATR Substrate Antibody (Cell Signaling Technology, #2851)

Anti-SDHA (Cell Signaling Technology, #5839)

Anti‐ 53BP1 (Cell Signaling Technology, #4937)

### Chemicals and assay kits

Agilent Seahorse XF Cell Mito Stress Test Kit (103015-100)

Alpha-Ketoglurate Colorimetric/Fluorometric Assay Kit (BioVision, K677-100)

Atractyloside (potassium salt) (Cayman, #14804)

ATP Colorimetric/Fluorometric Assay Kit (BioVision, K354-100)

B27 Supplement (50X), serum free (ThermoFisher Scientific, 17504044)

Carbonyl cyanide 4-(trifluoromethoxy)phenylhydrazone (FCCP) (Sigma, C2920-50MG)

Cytochrome Oxidase Activity Colorimetric Assay Kit (BioVision, K287-100)

Etoposide (Sigma, E1383-25MG)

GlutaMAX supplement (ThermoFisher Scientific, 35050061)

Gibson Assembly Master Mix (NEB, E2611)

High capacity cDNA Reverse Transcription kit (Applied Biosystems)

HyClone^TM^ Calf Serum (FisherScientific, SH3007203)

Hydrogen peroxide solution 30% (w/w) in water (Sigma, H1009-100ML)

Iscove’s modified Dulbecco medium (IMDM) (ThermoFisher Scientific, 12440053)

Isocitrate Colorimetric Assay Kit (Biovision, K656-100)

KU60019 (Selleckchem, S1570)

Lactate Colorimetric/Fluorometric Assay Kit (BioVision, K607-100)

L-Glutamic acid (Sigma, G1251-100G)

LIVE/DEAD^TM^ Viability/ Cytotoxicity Kit, for mammalian cells (L3224)

Lipofectamine 2000 (ThermoFisher Scientific, 11668019)

Lipofectamine LTX with Plus reagent (ThermoFisher Scientific, 15338100)

MEM Non-essential amino acid solution (100X) (ThermoFisher Scientific, 11140050)

Mitochondrial Complex I Activity Colorimetric Assay Kit (Biovision, K968-100)

Mitochondrial Complex III Activity Assay Kit (BioVision, K520-100)

MitoProfile^®^ Human Complex V Activity (Invitrogen, KHM2051)

Mitotracker Green FM (ThermoFisher Scientific, M7514)

N-Acetyl-L-cysteine (Sigma, A7250-25G)

Neurobasal medium (ThermoFisher Scientific, 21103049)

NE-PER^TM^ Nuclear and Cytoplasmic Extraction Reagents (ThermoFisher Scientific, 78833)

NU7441 (Tocris Bioscience, #3712)

Oligomycin (Sigma, 75351-5MG)

Opti-MEM^TM^ reduced serum mdium, GlutaMAX Supplement (ThermoFisher, 51985034)

Penicillin-Streptomycin (10,000U/mL) (ThermoFisher Scientific, 15140122) pCMBS (Clearsynth, CS-T-50790)

Pyruvate Colorimetric/Fluorometric Assay Kit (BioVision, K609-100)

Sodium Oxamate (Sigma, O2751-5G)

Succinate Dehydrogenase Activity Colorimetric Assay Kit (BioVision, K660-100)

Succinate (Succinic Acid) Colorimetric Assay Kit (BioVision, K649-100)

SYBR Green PCR Master Mix (Applied Biosystems)

Tetramethylrhodamine, Methyl Ester, Perchlorate (TMRM) (ThermoFisher Scientific, T668)

UK5099 (Merck-Calbiochem, CAS56396-35-1)

VE821 (Selleckchem, S8007)

2-Deoxy-D-glucose (Sigma, D8375-1G)

### Recombinant DNA

Laconic (Addgene, #44238)

pMX-3XFLAG-EGFP-OMP25 (Addgene, #83354)

pcDNA3.1(+)FLAG-HIS-ATM-wt (FLAG-ATM-WT) (Addgene, #31985)

pcDNA3.1(+)FLAG-HIS-ATM-kd (FLAG-ATM-KD) (Addgene, #31986)

pcDNA3.1(+)FLAG-HIS-ATM-C2991L (FLAG-ATM-C2991L) (in-house)

pEGFP-N1 (EGFP-vector) (Clontech)

pCMV6-EGFP-NRF1-WT (in-house)

pCMV6-EGFP-NRF1-NLSdel (in-house)

pCMV6-EGFP-NRF1-DIMdel (in house)

pCMV6-EGFP-NRF1-3XA (in house)

pCMV6-EGFP-NRF1-2XA (in house)

pCMV6-EGFP-NRF1-T259A (in house)

pCMV6-EGFP-NRF1-T259E (in house)

pCMV6-EGFP-NRF1-T201A (in house)

pCMV6-EGFP-NRF1-T287A (in house)

pCMV6-mCherry-NRF1-WT (in house)

pCMV6-mCherry-NRF1-T259A (in house)

pCMV6-mCherry-NRF1-T259E (in house)

pCMV6-Myc-FLAG-NRF1-WT (OriGene MR226726)

pCMV6-Myc‐ FLAG-NRF1-T201A (in house)

pCMV6-Myc‐ FLAG-NRF1-T259A (in house)

pCMV6-Myc‐ FLAG-NRF1-T259E (in house)

pCMV6-Myc‐ FLAG-NRF1-T287A (in house)

NRF1mitoGFP (A gift from Prof. Johan Tronstad)

Perceval High Range (Addgene, #49082)

Pyronic (Addgene, #51308)

Cyto-roGFP (Addgene, #49435)

### Sequence-Based Reagents

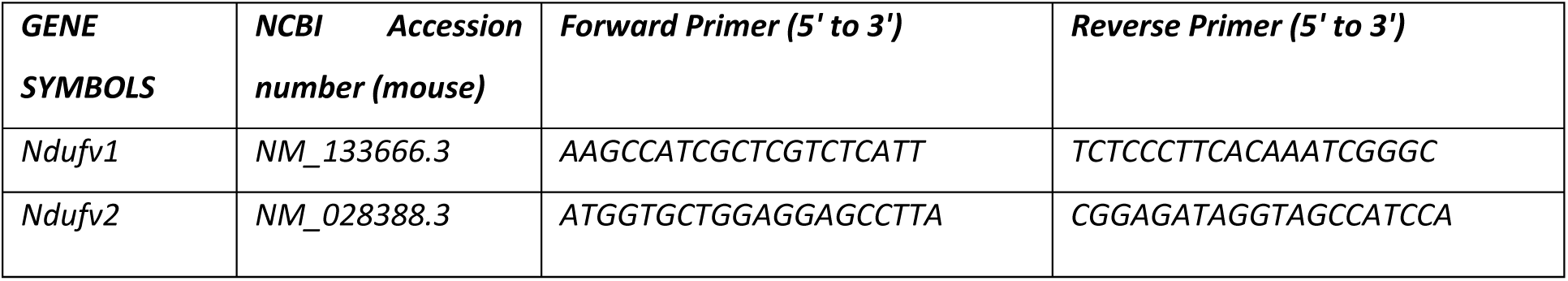
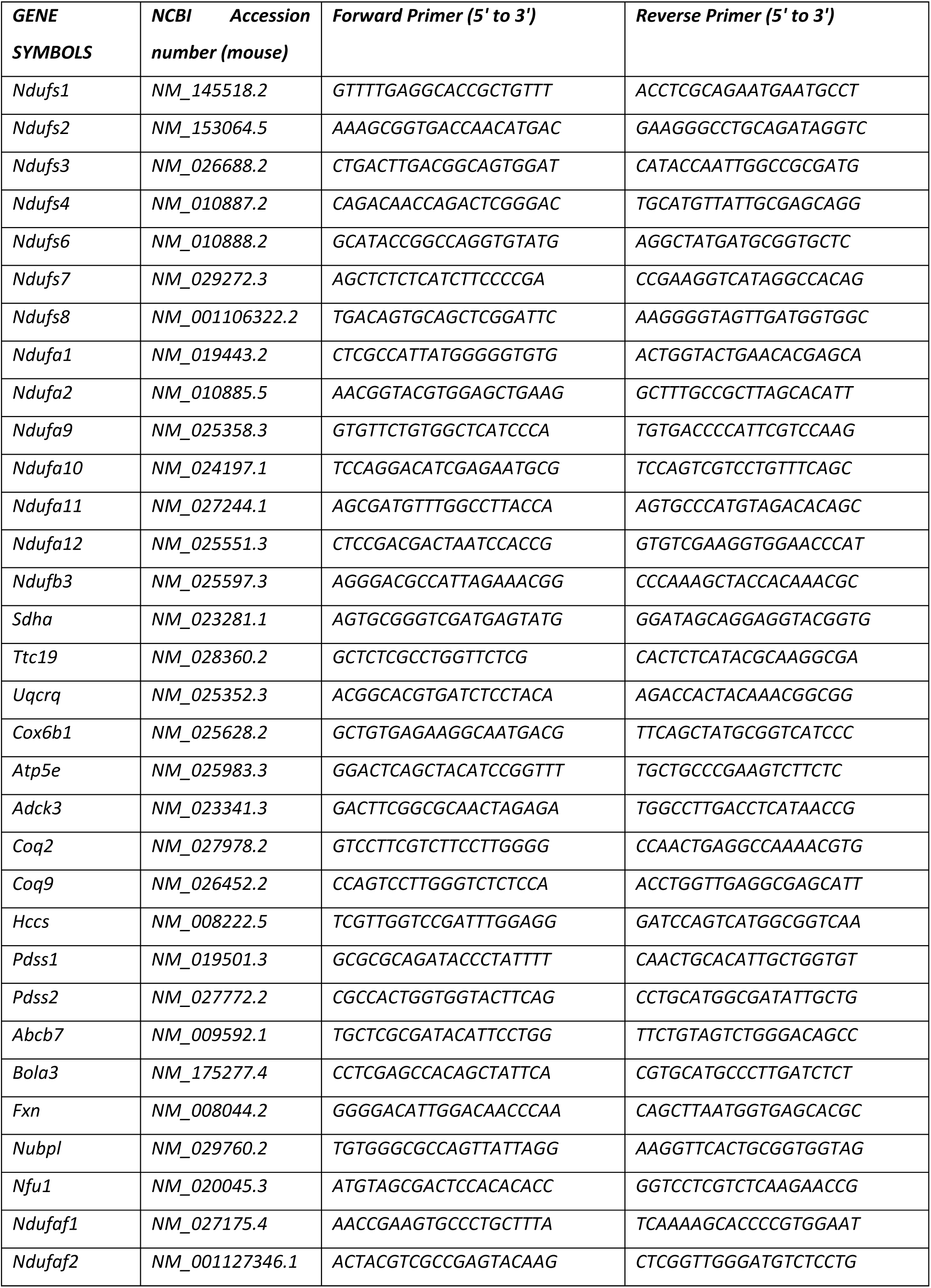
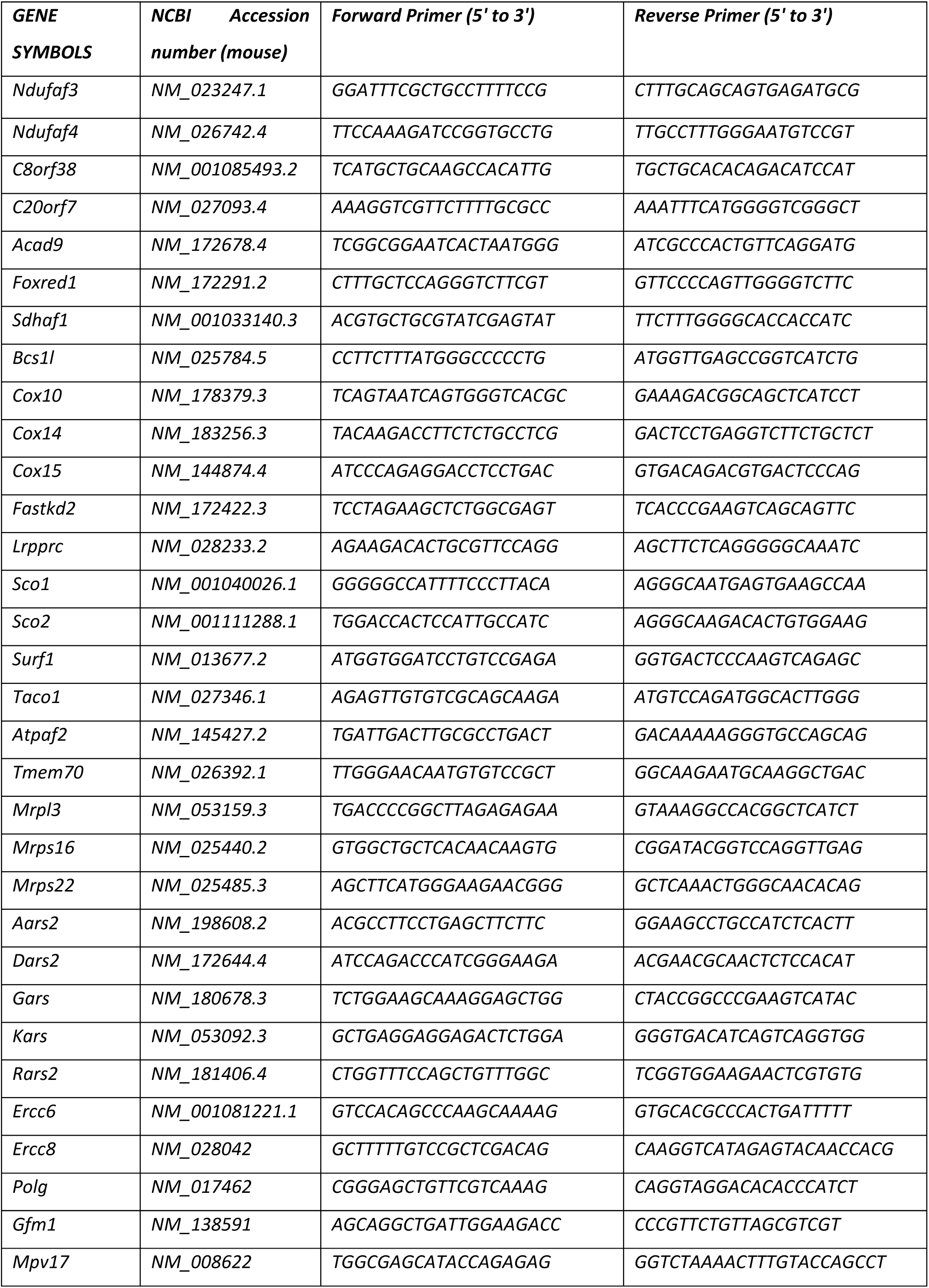
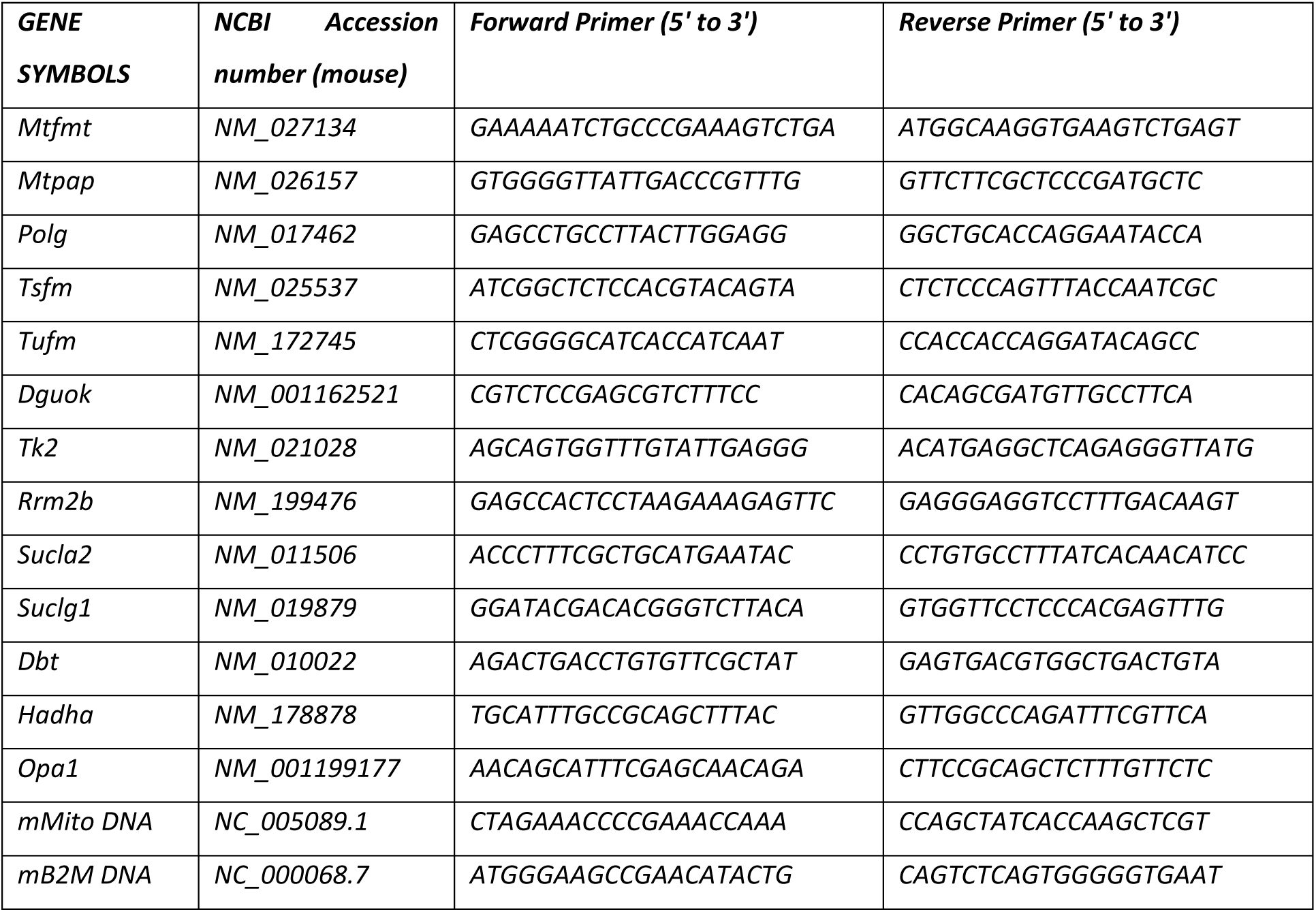
Primers for real time PCR

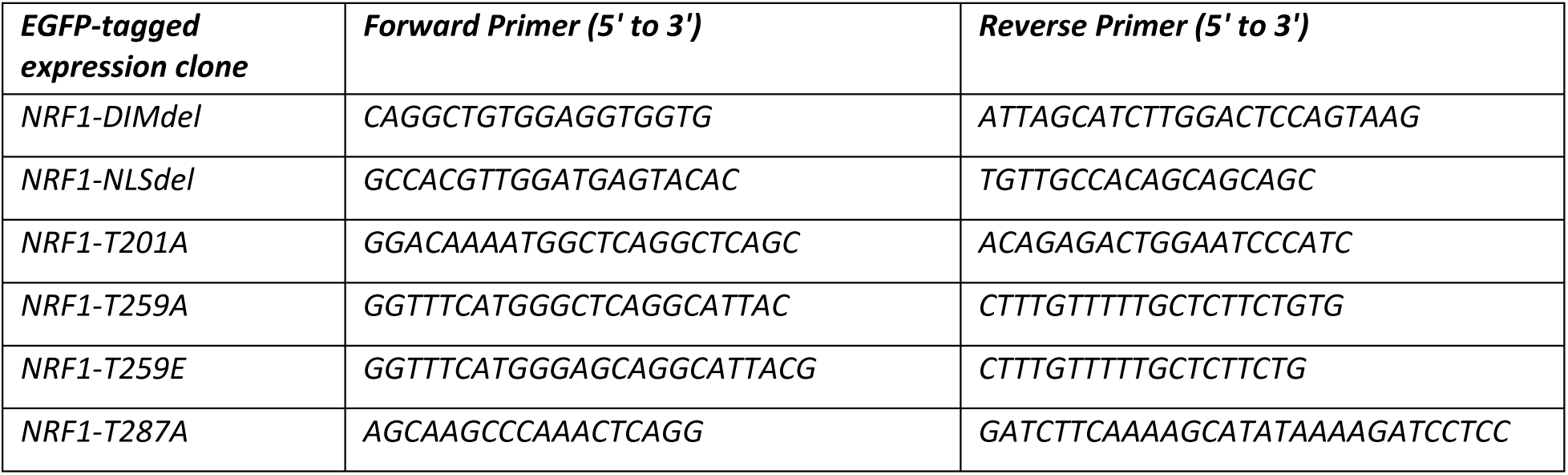
NRF1 mutant clones: Mutagenesis primers

### Software and Algorithms

GraphPad Prism 7

Molsoft ICM-Pro Browser

Pymol

Phyre2 http://www.sbg.bio.ic.ac.uk/phyre2/html/page.cgi?id=index

GEO2R https://www.ncbi.nlm.nih.gov/geo/geo2r

GSEA http://software.broadinstitute.org/gsea/index.jsp

GATHER http://changlab.uth.tmc.edu/gather/gather.py

MitoDB http://www.mitodb.com

MitoCarta2.0 https://www.broadinstitute.org/files/shared/metabolism/mitocarta/human.mitocarta2.0.html PSCAN http://159.149.160.88/pscan/

ZDOCK http://zdock.umassmed.edu

## Supplemental material

Fig. S1 shows impaired expression of OXPHOS genes in human A-T cerebellum. Fig. S2 shows loss of ATM impairs mitochondrial usage of pyruvate but enhanced lactate production in mouse neurons. Fig. S3 shows loss of ATM impairs ATP insufficiency response of human fibroblasts. Fig. S4 shows chronic insufficient ATP induces oxidative stress, activation of NRF1 and mitochondrial biogenesis. Table S1 shows raw data of multiple bioinformatics analyses.

## Acknowledgments

This work was financially supported by Research Grants Council, HKSAR (HKUST12/CRF/13G, GRF660813, GRF16101315, GRF16103317, GRF16100718 and AoE/M-05/12), the Offices of Provost, VPRG and Dean of Science, HKUST (VPRGO12SC02), and an RGC/HKUST initiation Grant (IGN16SC02), grant C6030-14E, and HKUST Institute for Advanced Study Junior Fellowship and Alzheimer’s Association Research Fellowship (AARF-17-531566) to HMC. MRS and RPH were supported by NIH R01 ES026057. We also acknowledge the staffs at the University Research Facility in Life Science (ULS) of Hong Kong Polytechnic University for their technical supports in experiments on the Agilent Seahorse platform.

### Author Contributions

HMC designed the experiments, performed the work, analyzed the data and wrote the manuscript. AC, XS and MS performed essential work, helped in the analysis of the data and helped in the writing of the experiments. RH and KH designed experiments, analyzed data and wrote the manuscript.

### Declaration of Interests

All authors declare no conflict of interests relate to the description and publication of the work described.

